# Thiopurines inhibit coronavirus Spike protein processing and incorporation into progeny virions

**DOI:** 10.1101/2022.03.10.483772

**Authors:** Eric S. Pringle, Brett A. Duguay, Maxwell P. Bui-Marinos, Rory P. Mulloy, Shelby L. Landreth, Krishna Swaroop Desireddy, Stacia M. Dolliver, Shan Ying, Taylor Caddell, Patrick D. Slaine, Stephen L. Bearne, Darryl Falzarano, Jennifer A. Corcoran, Denys A. Khaperskyy, Craig McCormick

**Author notes:** Co-corresponding authors: D.A.K., C.M. Co-lead authors (Eric S. Pringle, Brett A. Duguay).

## Abstract

There is an outstanding need for broadly acting antiviral drugs to combat emerging viral diseases. Here, we report that thiopurines inhibit the replication of the betacoronaviruses HCoV-OC43 and SARS-CoV-2, and to a lesser extent, the alphacoronavirus HCoV-229E. 6-Thioguanine (6-TG) disrupted early stages of infection, limiting synthesis of full-length and subgenomic HCoV RNAs. Furthermore, consistent with our previous report on the effects of thiopurines on influenza A virus glycoproteins, we observed that 6-TG inhibited accumulation of Spike glycoproteins from diverse HCoVs. Specifically, 6-TG treatment decreased the accumulation of Spike proteins and increased their electrophoretic mobility, consistent with Spike migration following the enzymatic removal of N-linked oligosaccharides with Peptide:N-glycosidase F (PNGaseF). SARS-CoV-2 virus-like particles (VLPs) harvested from 6-TG-treated cells were deficient in Spike. 6-TG treatment had a similar effect on lentiviruses pseudotyped with SARS-CoV-2 Spike; lentiviruses could be harvested from cell supernatants but were deficient in Spike and unable to infect human cells bearing ACE2 receptors. Together, these findings from complementary ectopic expression and infection models strongly indicate that defective Spike trafficking and processing is an outcome of 6-TG treatment. At low micromolar doses, the primary known mode of action of 6-TG is selective inhibition of the small GTPase Rac1. However, we observed that selective chemical inhibitors of the small GTPases Rac1, CDC42 and Rho had no effect on Spike processing and accumulation. The GTPase agonist ML099 countered the effects of 6-TG, suggesting that an unknown GTPase could be the relevant 6-TG-target protein involved in regulating Spike processing and accumulation. Overall, these findings provide important clues about the mechanism of action of a candidate antiviral that can broadly target HCoVs and suggest that small GTPases are promising targets for host-targeted antivirals.

**AUTHOR SUMMARY:** The COVID-19 pandemic has ignited efforts to repurpose existing drugs as safe and effective antivirals. Rather than directly inhibiting viral enzymes, host-targeted antivirals inhibit host cell processes to indirectly impede viral replication and/or stimulate antiviral responses. Here, we describe a new antiviral mechanism of action for the FDA-approved thiopurine 6-thioguanine. We demonstrate that this thiopurine is a pro-drug that must be metabolized by host enzymes to gain antiviral activity. We show that it can inhibit the replication of several human coronaviruses, including SARS-CoV-2, at least in part by interfering with the processing and accumulation of Spike glycoproteins, thereby impeding assembly of infectious progeny viruses. We provide evidence implicating host cell GTPase enzymes in the antiviral mechanism of action.

## INTRODUCTION

Coronaviruses (CoVs) are enveloped viruses with positive-sense, single-stranded RNA ((+)ssRNA) genomes. Seven human CoVs (HCoVs) are divided into two genera, the alphacoronaviruses (HCoV-NL63 and HCoV-229E) and betacoronaviruses (HCoV-OC43, HCoV- HKU1, MERS-CoV, SARS-CoV and SARS-CoV-2). Following entry into host cells, the (+)ssRNA CoV genome is immediately translated into polyproteins that are cleaved by viral cysteine proteases into 16 non-structural proteins (NSPs). These NSPs include the transmembrane nsp3-nsp4-nsp6 complex that generates a complex endoplasmic reticulum (ER)-derived reticulovesicular network that includes replication organelles known as double-membrane vesicles (DMVs) (1–8). In DMVs, the nsp3-nsp4-nsp6 complex provides a physical anchor for the viral replicase-transcriptase complex (RTC) that replicates the large viral genome and synthesizes subgenomic mRNAs (9, 10). These viral RNAs traverse a crown-shaped pore to exit the DMV (8). In the cytoplasm, viral mRNAs are translated into structural proteins; Nucleocapsid (N) is synthesized via free ribosomes in the cytoplasm, whereas mRNAs encoding envelope proteins are translated and processed in the ER and accumulate in the ER-Golgi-intermediate compartment (ERGIC). HCoV assembly takes place when full-length viral genomic RNA coated with N buds into the ERGIC to acquire the viral envelope and associated envelope proteins (11–13). These progeny viruses traverse the Golgi apparatus and *trans*-Golgi network (TGN), where envelope proteins receive additional post-translational modifications, and exit the cell via lysosomal exocytosis (14).

CoV envelope proteins are subject to extensive processing in the secretory pathway. The viral envelope proteins Spike, Envelope (E) and Matrix (M) that are found in all HCoVs, as well as Hemagglutinin-Esterase (H-E) found in only HCoV-OC43 and HCoV-HKU1, are synthesized in the ER as transmembrane proteins (15). CoV Spike proteins are large type I transmembrane proteins that are heavily N-glycosylated, which promotes proper folding, trimerization and trafficking (16). Subsequent proteolytic processing is required to activate these glycosylated Spike trimers, generating the S1 attachment subunit and liberating the mature amino-terminal fusion peptide of the S2 subunit, both of which are essential components of infectious CoV virions that can enter target cells (17, 18). Proteolytic activation of Spike can occur in the secretory pathway of infected cells or unprocessed Spike can be incorporated into virions and Spike processing can occur during attachment and entry into naïve cells in a process that involves multiple host serine and cysteine proteases. M is the most abundant CoV envelope protein and it serves as a scaffold for viral assembly. It is a multi-spanning membrane protein, with a small lumenal amino-terminus that is N-glycosylated, three transmembrane domains, and a large cytoplasmic carboxy-terminal domain known as the endodomain that governs interactions with the other structural proteins (Spike, E and N), as well as M-M interactions (19). E is a small type I transmembrane protein that forms pentameric complexes and functions as a viroporin, increasing the lumenal pH in the Golgi and protecting Spike from proteolysis (reviewed in (20)). The viroporin activity may support de- acidification of lysosomes and subsequent lysosomal exocytosis. M and E proteins both govern the intracellular trafficking and processing of Spike (19).

The striking remodeling of the ER to create replication organelles, as well as the reliance on the ER for synthesis and processing of viral glycoproteins, has prompted investigations into the effects of HCoV infection on ER stress and the unfolded protein response (UPR). Several CoVs have been shown to activate the UPR, including infectious bronchitis virus (IBV) (21, 22), mouse hepatitis virus (MHV) (23), transmissible gastroenteritis virus (TGEV) (24), HCoV-OC43 (25) and SARS-CoV (26–28). However, proximal UPR sensor activation does not always elicit downstream UPR transcription responses during CoV replication (29), which suggests complex regulation of the UPR by certain CoVs. Spike proteins from MHV (23), HCoV-HKU1 (27) and SARS-CoV (27, 30) are sufficient for UPR activation in cell culture. Additional transmembrane CoV proteins also induce the UPR, including SARS-CoV non-structural protein 6 (nsp6) (31), ORF3a (32) and ORF8ab (33) proteins. Despite these discoveries, our understanding of the regulation of the UPR in CoV infection remains incomplete, and little is known about UPR regulation in SARS-CoV-2 infection.

The emergence of SARS-CoV and MERS-CoV from animal reservoirs, followed by the current SARS-CoV-2 pandemic, has spurred efforts to discover and develop effective antivirals. These efforts have benefitted from drug repurposing strategies to identify antivirals amongst drugs that have already been approved for use in humans. We showed previously that thiopurines 6- thioguanine (6-TG) and 6-thioguanosine (6-TGo) selectively inhibit influenza A virus (IAV) replication by activating the UPR and interfering with the processing and accumulation of viral glycoproteins (34). Thiopurines have been in clinical use for over 50 years to treat inflammatory diseases and cancer and as immunomodulators. They include azathioprine, 6-mercaptopurine (6- MP) and 6-TG. 6-TG is orally bioavailable and on the World Health Organization’s List of Essential Medicines. It is used to treat hematological malignancies, including acute lymphoblastic leukemia (AML), acute lymphocytic leukemia (ALL) and chronic myeloid leukemia (CML), with the primary mechanism of action involving conversion into thioguanine deoxynucleotides and incorporation into cellular DNA, which preferentially kills rapidly dividing cancer cells (35, 36). By contrast, the anti-inflammatory effect of sub-cytotoxic doses of 6-TG for treating inflammatory bowel disease (IBD) is thought to be primarily due to inhibition of the small GTPase Rac1 (37, 38). This process requires 6-TG to be metabolized by the purine scavenging enzyme hypoxanthine phosphoribosyltransferase 1 (HPRT1) to yield 6-thioguanosine 5’-monophosphate (6-TGMP), which is then converted to 6-thioguanosine 5’-triphosphate (6-TGTP). 6-TGTP is the active metabolite that covalently modifies a reactive cysteine in the p-loop of Rac1. Subsequent hydrolysis of the GTPase-bound 6-TGTP adduct yields a trapped, inactive Rac1-6-TGDP product (37). These host-targeted properties of 6-TG, combined with our knowledge of how 6-TG inhibits IAV glycoprotein processing, motivated us to investigate whether 6-TG and related thiopurines might similarly interfere with CoV glycoproteins. Moreover, we reasoned that this kind of host-targeted antiviral activity might act synergistically with the previously reported direct-acting antiviral activity of 6-TG as an inhibitor of MERS-CoV, SARS-CoV and SARS-CoV-2 papain-like cysteine proteases PL^pro^ (nsp3) *in vitro* (39–42). Using ectopic expression and cell culture infection models, we discovered that 6-TG and 6-TGo inhibits replication of several CoVs including SARS-CoV-2, which correlates with disruption of Spike processing, as well as Spike accumulation and incorporation into progeny virions. Accumulation of viral subgenomic RNAs and full-length genomes is also inhibited by these thiopurines. Finally, we were able to link the antiviral mechanism of action to the primary known mode of action of 6-TG and related molecules as selective inhibitors of small GTPases (37, 43, 44), which could be overcome using a pan- GTPase agonist. Our findings extend the understanding of the mechanism of action of these candidate host-targeted antivirals and provide a means to study the impact of glycoprotein maturation on coronavirus biology.

## RESULTS

### 6-Thioguanine and 6-thioguanosine inhibit HCoV replication

We tested the effects of three thiopurines (**Fig. 1A**) on SARS-CoV-2 replication in Calu-3 lung adenocarcinoma cells. Treatment with 6-TG, the ribonucleoside 6-TGo, or the related thiopurine 6-MP, caused a striking 4-log decrease in the release of infectious virions after 48 h of infection, with little effect on cell viability [Selectivity Index (SI) 6-TG > 72,7, 6-Tgo > 81.6, 6-MP >153.8] (**Figs. 1B, 1C**). The sub-micromolar EC_50_ values that we observed for all three thiopurines in the Calu-3 infection model were consistent with previously reported values from other groups (42, 45). 6-TG was well tolerated in three other cell lines we used for HCoV infection models (HCT-8, Huh7.5, and primary hTERT-immortalized fibroblasts) (**Fig. 1D**). We tested the ability of 6-TG to inhibit the betacoronavirus HCoV-OC43 and the alphacoronavirus HCoV-229E and found that low micromolar doses of the ribonucleoside 6-TGo had similar inhibitory effects against HCoV-229E and HCoV-OC43 in their respective infection models, whereas 6-MP had no effect **(Figs. 1E, 1F)**. Because UPR stimulators thapsigargin (Tg) and tunicamycin (Tm) inhibit coronavirus replication (46–49) and our previous work demonstrated that 6-TG can inhibit glycosylation of influenza A virus (IAV) glycoproteins and stimulate ER stress (34), we included Tm as a positive control in these studies. In hTERT-BJ cells, 6-TG inhibited HCoV-OC43 replication at low micromolar doses, consistent with our observations in the HCT-8 cell infection model; however, HCoV-229E replication in hTERT-BJ cells was minimally affected across a range of sub-cytotoxic concentrations (**Figs. 1G, 1H**). Together, these findings suggest that 6-TG effectively inhibits replication of several coronaviruses.

**Figure 1.**
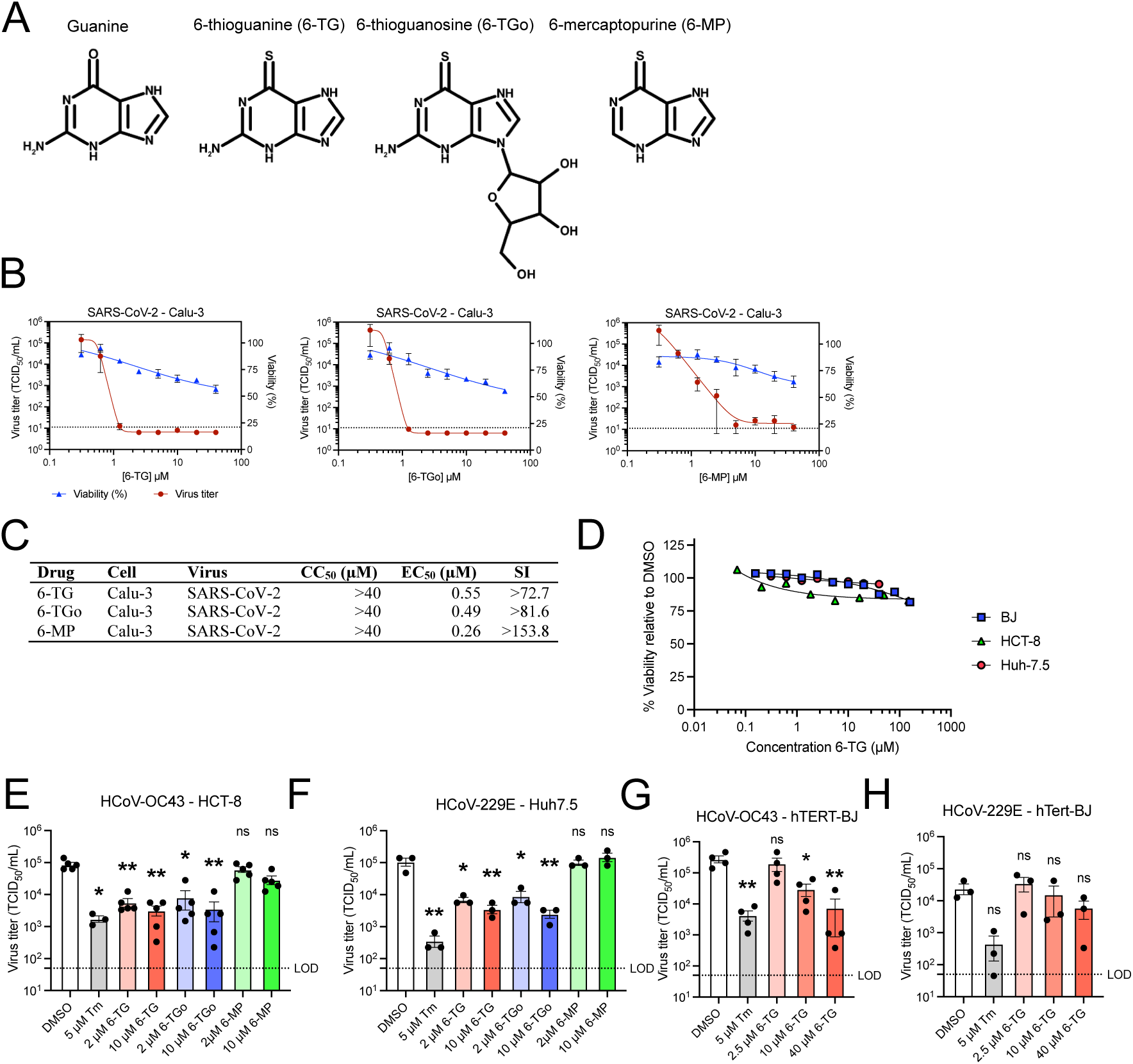
Thiopurines 6-thioguanine and 6-thioguanosine inhibit coronavirus replication. **(A)** Structures of thiopurines used this study in comparison to guanine. **(B)** Calu-3 cells were infected with SARS-CoV-2 at an MOI of 0.1 then treated with 6-thioguanine (6-TG), 6-thioguanosine (6- TGo), or 6-mercaptopurine (6-MP). Supernatant was harvested after 48 h, and stored at -80∞C until titering on Vero’76 cells. Mock-infected cells were similarly treated with 6-TG, 6-TGo, 6- MP, or DMSO vehicle control for 48 h before testing cell viability with CellTiter 96 AQueous One (n=3 ± SEM). Dotted line indicated Limit of Detection. **(C)** Summary table of 50% Cytoxic Concentration (CC_50_), 50% Effective Concentration (EC_50_), and Selectivity Index (SI) calculated for (A-C). **(D)** Alamar Blue cell viability assay of hTert-BJ, HCT-8, and Huh-7.5 cells treated with 6-TG (n=3±SEM). **(E)** TCID assays for HCoV-OC43 infected HCT-8 cells and **(F)** HCoV-229E infected Huh-7.5. Cells were infected with an MOI of 0.1 then treated with 6-TG, 6-TGo, 6-MP, or DMSO (n=3 ± SEM, statistical significance was determined by one-way ANOVA). **(F)** hTERT- BJ cells were infected with HCoV-OC43 or HCoV-229E at an MOI of 0.1 and treated with 6-TG, Tm, or DMSO. Supernatant was harvested after 23 h and stored at -80∞C before titering on BHK- 21 or Huh7.5 (n=3-4 ± SEM, statistical significance was determined by one-way ANOVA). **(G)** hTERT-BJ cells were infected with HCoV-OC43 or **(H)** HCoV-229E at an MOI of 0.1 and treated with 6-TG, Tm, or DMSO. Supernatant was harvested after 23 h and stored at -80∞C before titering on BHK-21 or Huh7.5 (n=3-4 ± SEM, statistical significance was determined by one-way ANOVA). LoD = Limit of Detection for virus titer. (*, p<0.05; **, p<0.01; ns, non-significant)

### 6-Thioguanine impedes viral genome replication and subgenomic RNA synthesis

Next, we investigated which stages of virus replication were affected by 6-TG treatment. We performed multi-round HCoV-OC43 infections of 293T cells at an MOI of 0.1. Infected cells were treated with 10 µM 6-TG or DMSO immediately after infection, and cell lysates and supernatants were harvested at regular intervals for the next 48 hours. 6-TG treatment caused a ∼40-fold reduction in viral titer during the first day of infection. Virion release from control cells plateaued at 24 hours post-infection, whereas peak release of infectious virions from 6-TG-treated cells occurred much later at 48 hours post-infection **(Fig. 2A)**. Similarly, measuring viral particle release by RT-qPCR amplification of extracellular viral genomes revealed a ∼20-fold reduction in virus titer **(Fig. 2B)** throughout the first day of infection.

**Figure 2.**
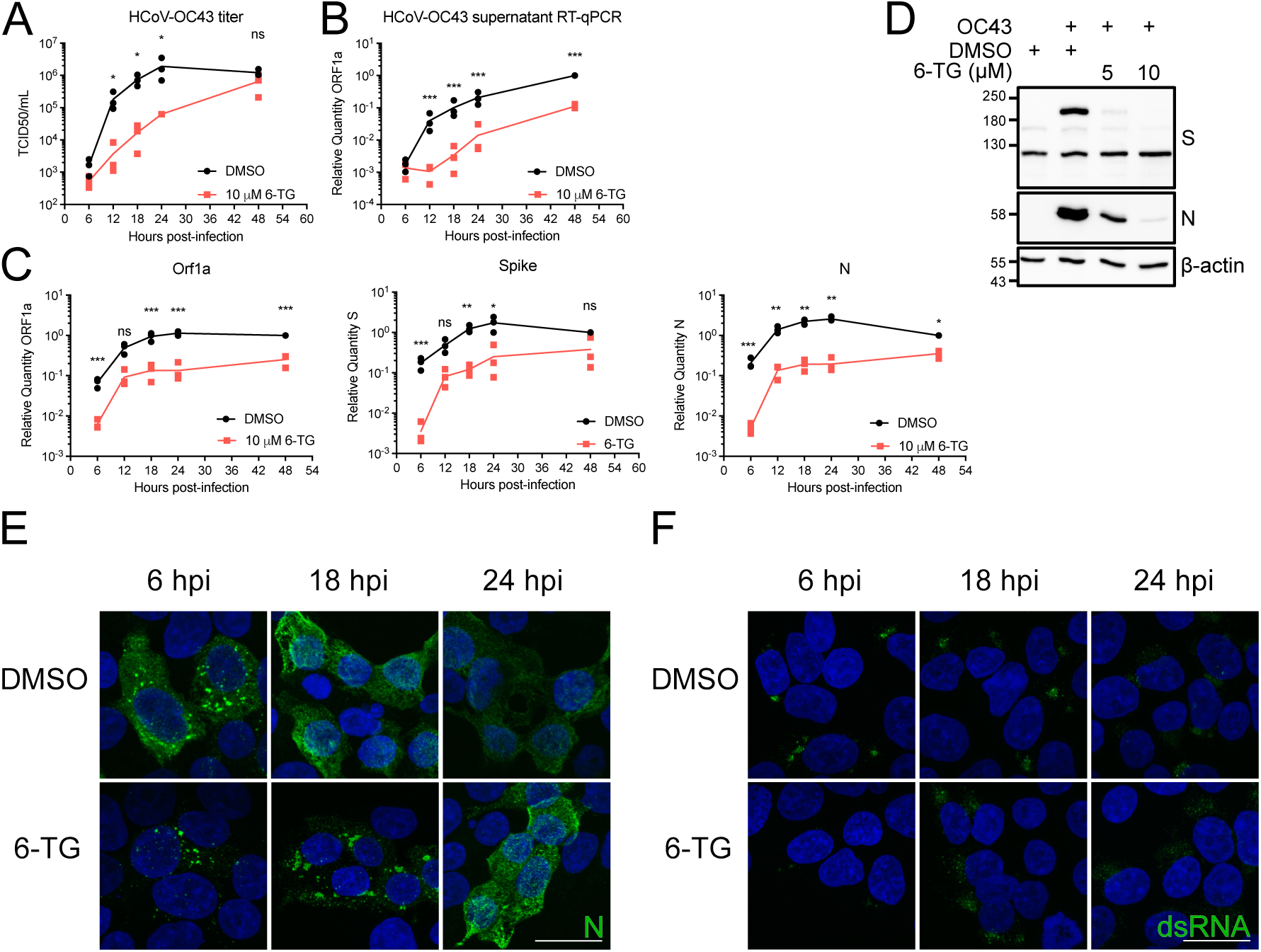
6-TG reduces CoV protein and mRNA accumulation. **(A-C)** 293T cells were infected with HCoV-OC43 at an MOI of 0.1 then treated with 10 µM 6-TG or DMSO vehicle control. Supernatant and total RNA from the cells were harvested at the times indicated and stored at - 80∞C until **(A)** titering on BHK-21 or **(B)** quantification of viral genomes by RT-qPCR. Cell monolayers were harvested for **(C)** RT-qPCR analysis of viral RNA accumulation (n=3 statistical significance was determined by paired ratio t test; *, p<0.05; **, p<0.01; ns, non-significant).**(D)** 293T cells were infected as in (A) then treated with 5 or 10 µM 6-TG or DMSO. Lysates were harvested 24h after infection and were probed by western blotting as indicated. **(E)** 293T cells were grown on coverslips then infected with HCoV-OC43 and treated with 6-TG as in (A), then fixed and immunostained for N or **(F)** dsRNA. Nuclei are counterstained with Hoechst. Scale bar = 20 µm.

Next, we investigated the ability of 6-TG to disrupt viral RNA synthesis in infected cells. Putative full-length RNA, as detected by primers targeting *ORF1a*, was reduced by ∼10-fold throughout most stages of infection. Similar decreases were detected using primers targeting the coding region of *Spike* and *N* genes after 6 h post-infection, with much larger 40-fold differences at 6 h post-infection **(Fig. 2C)**. Accumulation of viral proteins N and Spike was also inhibited (Fig. 2D). These disruptions in viral RNA accumulation correlated with changes in the localization of N, the viral nucleoprotein. Immunostaining HCoV-OC43 infected cells with an anti-N antibody revealed a punctate staining pattern at 6 hours post-infection **(Fig. 2E)**. By 24 h post-infection, N immunostaining was less punctate and more peripheral. The N immunostaining was markedly altered in 6-TG-treated infected cells, with brighter punctate staining patterns at 6 hours post-infection and large puncta still visible at 24 h post-infection. The formation of replication compartments was also delayed in 6-TG treated cells, as determined by immunostaining for dsRNA **(Fig. 2F)**; Specifically, we observed dsRNA signal at 6 hpi, which correlated with decreased viral RNA detected by RT-qPCR **(Fig. 2C)**. Together, these findings indicate that 6-TG reduces the synthesis of viral RNAs required to support efficient HCoV replication.

In our recent study, the antiviral mechanism of action of 6-TG and 6-TGo against IAVs involved activation of the UPR and interference with the processing and accumulation of viral glycoproteins. In these experiments, IAV infection of vehicle-treated A549 cells revealed very little detectable UPR late in infection at 24 h post-infection, whereas 6-TG stimulated all three arms of the UPR, with transactivation of XBP1s target gene *Erdj4*, ATF6-N target genes *BiP* and *HERPUD1* and ATF4 target gene *CHOP* (34). Thus, thiopurines markedly altered the ER proteostasis landscape to the detriment of IAV replication. Notably, many HCoVs have been shown to activate the UPR, including infectious bronchitis virus (IBV) (21, 22), mouse hepatitis virus (MHV) (23), transmissible gastroenteritis virus (TGEV) (24), human coronavirus (HCoV)- OC43 (25) and SARS-CoV (26–28). Here we show that HCoV-OC43 infection caused induction of *Xbp1* transcript, potent accumulation of spliced Xbp1, and transactivation of XBP1s target genes *Erdj4* and *Edem1*, providing strong evidence for activation of the IRE1 arms of the UPR **(Figs. S1A, S1B)**. Regulation of the PERK/ATF4 arm of the UPR may be more complex, as HCoV- OC43 infection caused accumulation of *ATF3* mRNA but no change in *CHOP*, even though they are both encoded by ATF4 target genes. Similarly, we observed increased expression of ATF6-N target genes *HERPUD1* and *Xbp1* but no increases in BiP mRNA or protein **(Figs. S1A, S1B)**. Thus, HCoV-OC43 interactions with the UPR appear to be complex and marked by strong stimulation of IRE1, which is consistent with observations from others (25). In marked contrast to our IAV studies, 6-TG treatment generally delayed or suppressed the downstream UPR transcriptional response **(Fig. S1A)**, which could be attributed to 6-TG-mediated delays in viral replication and the accumulation of viral RNAs and proteins **(Fig. 2)**. Finally, it should be noted that UPR transcripts accumulated to high levels in vehicle-treated cells despite the fact that HCoV- OC43, like all HcoVs, has mechanisms to potently shut off host antiviral gene expression (25). Using ribopuromycinylation assays, we demonstrated that HCoV-OC43 greatly reduced translation of host mRNAs at the expense of viral mRNAs encoding a few abundant viral proteins **(Fig. S1C, lane 4)**. 6-TG treatment had no effect on global rates of translation initiation in HCoV- OC43 infected cells, with the only changes evident being the apparent loss of some abundant viral proteins **(Fig. S1C, lane 6)**. Together, these observations suggested that 6-TG inhibition of HCoV- OC43 infection limits the strong activation of IRE1 and accumulation of XBP1s target genes, and that even though 6-TG interferes with synthesis of viral full-length genomic RNAs and subgenomic RNAs, host shutoff is unperturbed.

### 6-Thioguanine inhibits Spike processing and accumulation

We previously reported that 6-TG and 6-TGo inhibited the processing and accumulation of IAV glycoproteins hemagglutinin (HA) and neuraminidase (NA) (34). Here, we observed a similar defect in SARS-CoV-2 infected cells; lysates harvested from SARS-CoV-2-infected Calu-3 cells at 48 h post-infection showed a sharp decrease in all four structural proteins following 6-TG treatment **(Fig. 3A)**, consistent with a sub- micromolar EC50 discussed above **(Fig. 1B)**. Treatment of SARS-CoV-2-infected Huh-7.5 cells with 10 μM 6-TG also reduced the accumulation of viral structural proteins, which was especially evident for Spike **(Fig. 3B)**. These findings led us to further investigate the effects of 6-TG on SARS-CoV-2 structural proteins. Transfection of 293T cells with plasmids encoding each SARS-CoV-2 structural protein revealed that 6-TG had no effect on E or N accumulation, whereas Spike processing was noticeably inhibited (**Fig. 3C**). We also observed that 6-TG treatment caused a loss of a slow-migrating species of M **(Fig. 3C)**, which likely represented the glycosylated form of the protein that has been previously reported for other HCoVs (50). Thus, it appears that amongst the SARS-CoV-2 structural proteins, only the glycosylated Spike and M proteins are altered by 6-TG treatment when expressed independently in 293T cells.

**Figure 3.**
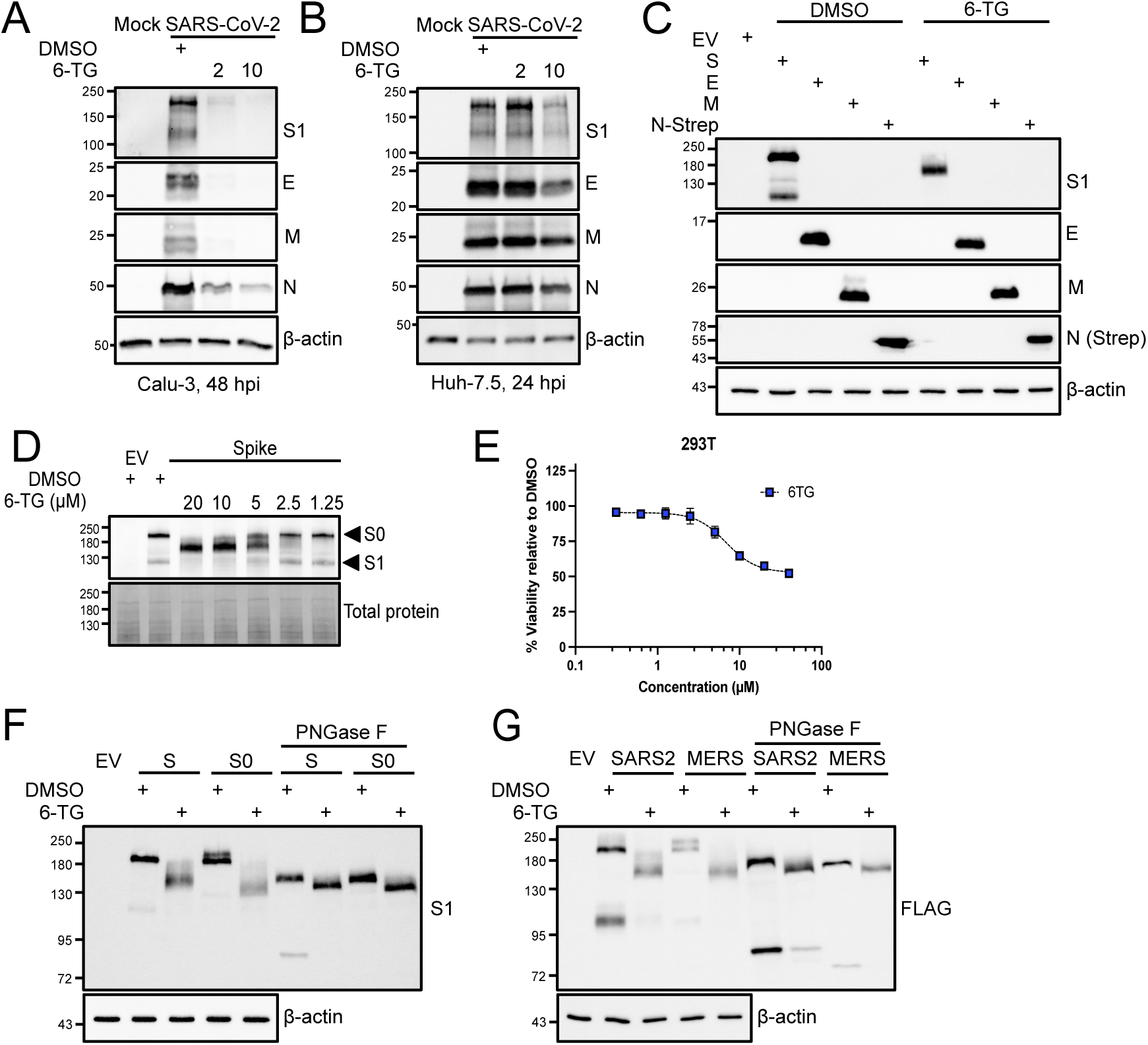
6-TG inhibits Spike glycosylation and cleavage. **(A)** Calu-3 cells were infected with SARS-CoV-2 at an MOI of 7, then treated with 6-TG or DMSO. Lysates were harvested 48 h after infection and were probed by western blotting as indicated. Concentrations in legend are in µM. **(B)** Huh-7.5 cells were infected with SARS-CoV-2 at an MOI of 5, then treated with 6-TG or DMSO. Lysates were harvested 48 h after infection and were probed by western blotting as indicated. Concentrations in legend are in µM. **(C)** 293T cells were transfected with SARS-CoV- 2 S, M, Envelope (E), Nucleoprotein (N) with a C-terminal 2xStrep tag, or EV then treated with 10 µM 6-TG or DMSO vehicle control. Lysates were harvested 24 h after transfection and probed by western blotting as indicated. **(D)** 293T cells were transfected with SARS-CoV-2 Spike or EV then treated with 6-TG or DMSO vehicle control. Lysates were harvested 24 h after transfection and were probed by western blotting as indicated. Concentrations in legend are in µM. **(E)** Alamar Blue cell viability assay of 6-TG treated 293T cells (n=3±SEM). **(F)** 293T cells were transfected with SARS-CoV-2 Spike (S), a Spike mutant lacking the Furin cleavage site between S1 and S2 (S0), or EV then treated with 10 µM 6-TG or DMSO vehicle control. Lysates were harvested 24 h after transfection, treated with PNGase F to remove N-linked glycans, and probed by western blotting as indicated. **(G)** as in (C) except 293T cells were transfected with C-terminal FLAG- tagged Spike (S-FL) or HCoV-MERS Spike (MERS).

To further explore the effects of 6-TG on Spike, we ectopically expressed Spike, then treated the cells with a range of sub-cytotoxic 6-TG concentrations before harvesting lysates and processing for immunoblotting **(Figs. 3D, 3E)**. Spike appeared as two bands when immunoblots are probed with an anti-S1 receptor binding domain (RBD) antibody, a high-molecular weight (MW) precursor S0 band (∼203 kDa) and a lower molecular weight band representing the protease-cleaved S1 attachment protein (∼96 kDa). Increasing concentrations of 6-TG once again led to the accumulation of a prominent single band at an unexpected MW (∼137 kDa) **(Fig. 3D)**. Because 6- TG treatment caused Spike to migrate faster during SDS-PAGE than expected, we hypothesized that the lower MW species observed only in 6-TG treated samples could arise from a glycosylation defect. We treated lysates from Spike-expressing 293T cells with Peptide-*N*-Glycosidase F (PNGase F), an amidase that cleaves the bond between N-acetylglucosamine (GlcNAc) and asparagine, effectively removing all N-linked glycans from the polypeptide chain (unless the core GlcNAc is modified by *α*-1,3-fucose) (51). PNGase F treatment caused Spike from the DMSO-treated cells to migrate similarly to Spike from 6-TG-treated cells **(Fig. 3F)**. We confirmed the identity of the high MW Spike band as the uncleaved S0 precursor protein by generating and testing a mutant Spike protein in which the basic residues in the furin cleavage site joining S1 and S2 were substituted with alanines, yielding a protease cleavage-resistant Spike protein. Cells transfected with this S0 construct and treated with 6-TG prior to lysis revealed that the S0 construct was also sensitive to PNGaseF treatment; the 6-TG/PNGaseF-treated S0 migrated similarly to 6-TG/PNGaseF-treated WT Spike by SDS-PAGE. Several studies have mapped O-linked glycans on SARS-CoV-2 Spike (52–54) and these modifications appear to follow the “O-follow-N rule” whereby N-glycosylation is followed by O-glycosylation on nearby residues (52). Thus, the apparent differences in molecular weight between vehicle/PNGaseF-treated Spike and 6- TG/PNGaseF-treated Spike could stem from residual O-linked glycans on Spike that are absent in the 6-TG-treated proteins. These findings were corroborated using carboxy-terminal FLAG-tagged Spike proteins from SARS-CoV-2 and MERS-CoV that also displayed a fast-migrating, under- glycosylated S0 species and loss of the S2 subunit following 6-TG treatment **(Fig. 3G)**. Taken together, these results suggest that 6-TG inhibits normal glycosylation and processing of Spike.

A failure of Spike to migrate efficiently from the ER to the Golgi could inhibit glycan maturation and cleavage by furin. In our previous work, we observed that 6-TG treatment did not reduce levels of Class I HLA proteins at the cell surface (34), which we interpreted as evidence that 6-TG did not globally affect processing and secretion of host cell glycoproteins. Here, we tested the role of 6-TG on the secretory pathway by using a *Gaussia* luciferase assay in which the enzyme depends on a native secretion signal to be secreted (55). As a control we used the Arf1- inhibitor brefeldin A (BFA) which potently inhibits protein trafficking from the ER to the Golgi (56, 57). BFA is toxic to cells at micromolar doses, but we could still detect secretory pathway inhibition at 250 nM for 20 h without overt cytotoxicity **(Figs. S2A, S2B)**. A short-term, 6 h treatment with 6-TG (10 µM) had little effect on *Gaussia* luciferase secretion, whereas treatment with BFA (10 µM) greatly decreased secretion. We tested a longer 20 h treatment of 6-TG where we co-transfected Firefly luciferase (Luc2) with *Gaussia* luciferase to help control for the inhibitory effects of 6-TG on 293T cell proliferation. This extended 6-TG treatment caused a slight loss of detectable *Gaussia* luciferase activity in the cell supernatant, but once again low-dose treatment of BFA greatly inhibited secretion and increased intracellular retention of *Gaussia* luciferase (**Fig. S2C**). These experiments strongly corroborated our prior observations that 6-TG treatment does not globally disrupt the secretory pathway. Because of our previous observations of 6-TG-mediated defects in Spike glycosylation, processing and trafficking, we decided to test whether low-dose, long-term treatment with BFA would be sufficient to recapitulate the effects of 6-TG treatment on Spike. However, unlike 6-TG, the secretion inhibitor BFA did not cause any changes in Spike accumulation or processing, as indicated by identical electrophoretic mobility of Spike on immunoblots compared to vehicle control **(Fig. S2D).**

To directly monitor Spike protein secretion, we used flow cytometry and surface staining of unpermeablized 293T cells transfected with Spike. We co-transfected cells with an EGFP plasmid and gated on EGFP+ cells to measure changes in Spike accumulation. 6-TG reduced surface expression of Spike by ∼50% resulting in fewer Spike-positive cells and reduced mean fluorescence intensity. In these experiments, we also tested a Spike protein in which the 19 C-terminal residues were removed (Spike-Δ19); this widely-used strategy removes an ER-retrieval signal and enhances incorporation of Spike into lentivirus particles (58–60). We found a similar ∼50% reduction in trafficking and surface expression of Spike-Δ19 **(Fig. S2E)**. Together, these experiments indicate that 6TG inhibits normal trafficking of Spike to the cell surface but does not broadly inhibit secretion.

### 6-Thioguanine causes a coronavirus assembly defect

It has been shown that co-transfection of plasmids expressing each of the four SARS-CoV-2 structural proteins allows for secretion of virus-like particles (VLPs) (19), which is concordant with earlier studies with other CoV structural proteins (61). However, we observed that when Spike, E, M and N expression vectors were co-transfected into cells, 6-TG treatment not only altered the accumulation and processing of Spike, but also dramatically reduced accumulation of E and M proteins (**Fig. 4A**). Cell supernatants were collected, filtered and pelleted in a sucrose cushion to isolate VLPs, which were then characterized by immunoblotting. We observed a dramatic reduction in VLP production from 6-TG-treated 293T cells, and these residual VLPs completely lacked Spike (**Fig. 4B**). These findings indicate that 6- TG not only prevents Spike processing and accumulation, but also its incorporation into VLPs. We then co-transfected cells again with only three of the four Spike, E, M, and N encoding plasmids to better understand mechanisms of defective assembly. Our main conclusions are: 1) 6- TG treatment of cells expressing all structural proteins causes a strong Spike-dependent decrease in E expression, 2) in the absence of M the 6-TG-mediated Spike glycosylation defect is more pronounced, and 3) N is required for accumulation of E **(Fig 4C)**. To determine if Spike from 6- TG treated cells was still functional, we employed a lentivirus pseudotyping method using Spike- Δ19, which is widely used to measure neutralizing antibody titers to SARS-CoV-2 (58, 62, 63). We performed immunoblots on pseudotyped viruses (PVs) recovered from three independent preparations that we purified by ultracentrifugation through a 20% sucrose cushion. Unlike our previous immunoblots of ectopically expressed Spike from transfected cells, we observed that most of the Spike proteins incorporated into the PVs were fully cleaved into the prominent S1 band **(Fig. 4D)**. However, 6-TG treatment led to a dramatic loss of Spike S0 and S1 species from the purified PVs, compared to the smaller decrease observed for HIV p24 levels in PV cores. We quantified PV yield from these cells by amplifying lentiviral genomes via RT-qPCR from filtered, unconcentrated cell supernatants. 6-TG treatment did not significantly reduce the quantity of capsid-protected genomes released from cells compared to vehicle control **(Fig. 4E)**. To determine the effects of 6-TG on PV infectivity, we used purified PVs to infect HEK293A cells that stably express the ACE2 receptor and measured luciferase production in the recipient cells as a measure of successful lentiviral infection. Consistent with deficient Spike incorporation into PVs, we observed that 6-TG treatment reduced PV infectivity by over 50-fold compared to PVs derived from vehicle-treated cells **(Fig. 4F)**. By contrast, PVs derived from vehicle control- and 6-TG-treated cells were equally competent to infect ACE2-deficient HEK293A cells at low levels. This suggests that 6-TG treatment not only inhibits Spike glycosylation and processing, but it also inhibits Spike trafficking and incorporation into lentiviral particles. We next investigated whether these 6-TG-mediated assembly defects altered the morphology of HCoV particles. We concentrated supernatant from HCoV-OC43 infected 293A cells by ultracentrifugation and imaged virions by transmission electron microscopy using negative staining. We observed fewer viral particles in the 6-TG treated samples, and while these particles were of similar size, we noted fewer particles with clearly discernible Spikes **(Fig. 4G)**. Taken together, the strong effects of 6-TG on Spike in transiently transfected cells, heterologous lentivirus production and infection, and authentic HCoV infections, provide evidence for an additional antiviral mode of action for 6-TG beyond PLpro/nsp3 inhibition or defects in viral RNA synthesis.

**Figure 4.**
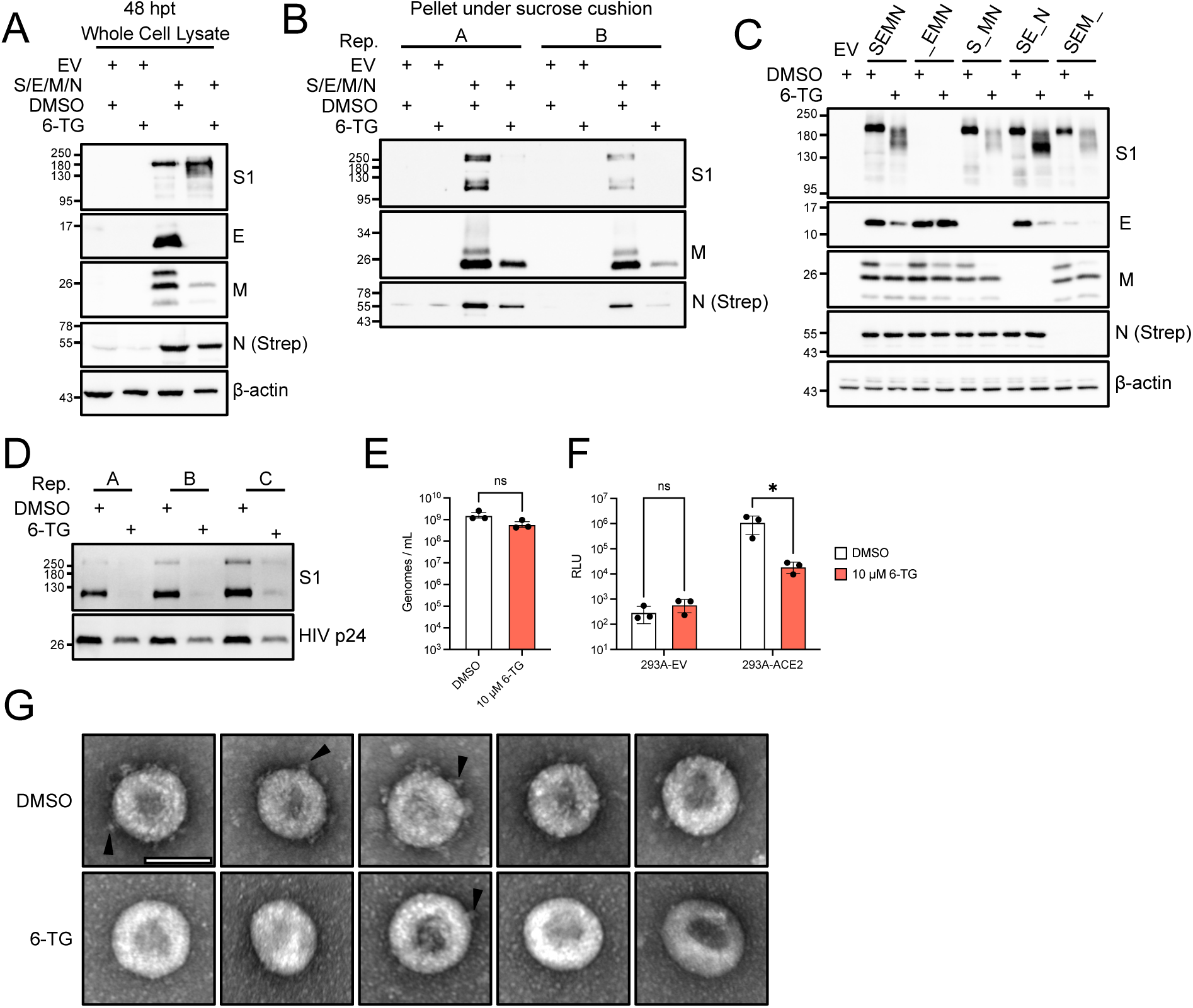
6-TG stimulates an assembly defect and inhibits Spike virion incorporation. **(A)** 293T cells were transfected with equal quantities of SARS-CoV-2 S, M, E, and N plasmids or empty vector (EV) then treated with 10 µM 6-TG or DMSO vehicle control. Lysates were harvested 48 h after transfection and probed by western blotting as indicated. **(B)** Virus-like particles from supernatants of cells transfected in (B) were concentrated by ultracentrifugation through a 20% sucrose cushion, resuspended in PBS, then harvested with an equal volume of 2x Laemmli buffer. Samples from two independent lentivirus preparations were probed by western blotting as indicated. **(C)** 293T cells were transfected with S, E, M, and N in a 1:2:2:1 ratio, substituting one of the structural proteins for EV as indicated, treated with 6-TG or DMSO then processed as in (A). **(D)** SARS-CoV-2 Spike pseudotyped, luciferase-expressing lentivirus particles were concentrated by ultracentrifugation through a 20% sucrose cushion, resuspended in PBS, then harvested with an equal volume of 2x Laemmli buffer. Samples from three independent lentivirus preparations were probed by western blotting as indicated. **(E)** Genomes from three independent lentivirus preparations were quantified by RT-qPCR (n=3, statistical significance was determined by paired t-test; ns, non-significant). **(F)** 293A cells stably expressing ACE2 or empty vector control were transduced with lentivirus from three independent preparations. After 24 h, lysates were harvested and measured for luciferase activity (n=3, statistical significance was determined by two-way ANOVA; *, p<0.05; ns, non-significant). **(G)** 293A cells were infected HCoV-OC43 at an MOI of 0.1 =, then treated with 6-TG or DMSO. Supernatants were concentrated as in (C), before virions were fixed and imaged by TEM with negative staining. Five virions from both 6-TG- and DMSO-treated samples are shown at 150,000 X magnification. Scale bar = 100 nm. Arrowhead indicates examples of Spike protein extending from virion.

### 6-Thioguanine must be processed by HPRT1 to inhibit Spike processing and coronavirus replication

We observed comparable antiviral activities of 6-TG and its ribose-conjugated analogue 6-TGo throughout these studies. Metabolism of 6-TG is initiated by the purine scavenging enzyme hypoxanthine phosphoribosyltransferase 1 (HPRT1) to yield 6-thioguanosine 5’-monophosphate (6-TGMP) **(Fig. 5A)**. 6-TGMP can then be converted to 6-thioguanosine 5’- triphosphate (6-TGTP), the active form of the molecule that has been shown to form a covalent bond with a reactive cysteine in the p-loop of Rac1; subsequent hydrolysis of Rac1-6-TGTP to a diphosphate form renders the GTPase inactive (37). Considering this information, we next sought to determine whether conversion of 6-TG to a ribonucleoside was required for its antiviral activity. First, we synthesized N9-methyl 6-thioguanine (6-TG-Me), which prevents conjugation to a ribose sugar due to the methyl group at N9 of the purine ring and is predicted to be resistant to conversion into 6-TGMP **(Fig. 5A)**. 6-TG-Me displayed low cytotoxicity and no antiviral activity against HCoV-OC43 across a broad range of concentrations **(Fig. 5B)**. Likewise, 6-TG-Me had no effect on SARS-CoV-2 Spike processing across a range of concentrations between 5 μM and 20 μM **(Fig. 5C)**. To corroborate this finding, we used RNA interference to silence expression of *HPRT1* in 293T cells prior to infection with HCoV-OC43 and treatment with 6-TG or vehicle control. RT-qPCR confirmed effective silencing of *HPRT1* **(Fig. 5D**), which had no effect on HCoV-OC43 infection in vehicle control-treated cells **(Fig. 5E)**. By contrast, *HPRT1* silencing significantly restored viral replication in the presence of 6-TG, although replication did not return to the levels in control cells. Together, these findings indicate that 6-TG is a pro-drug that must be metabolized by HPRT1 to convert it to an active form.

**Figure 5.**
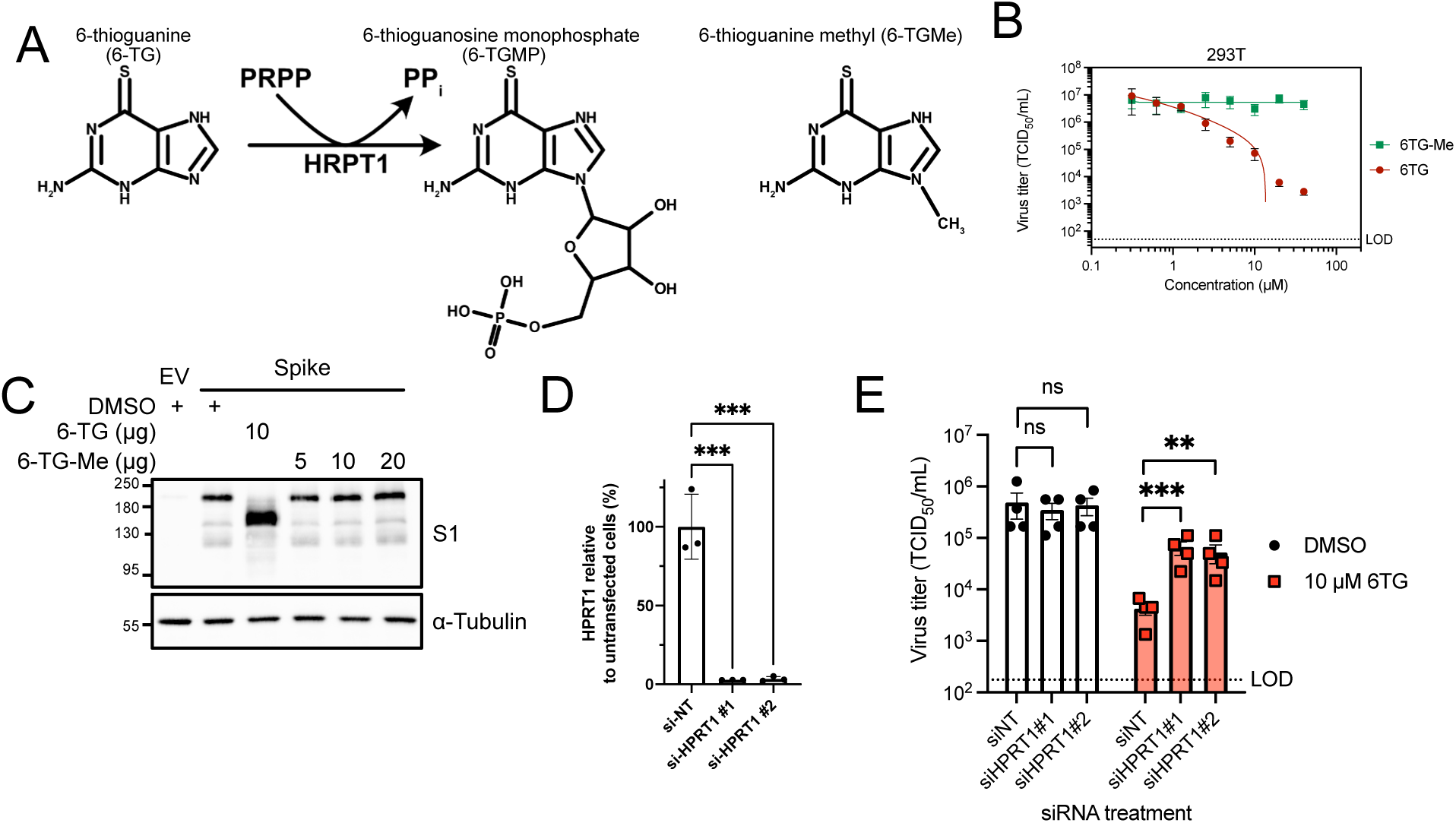
6-TG inhibits Spike maturation through GTPase inhibition. **(A)** 6-TG is conjugated to phosphoribosyl diphosphate (PRPP) by HPRT1 to form 6-TG monophosphate. 6-TG methylated at the N9 nitrogen was designed to be resistant to processing by HPRT1. **(B)** 293T cells were infected with HCoV-OC43 at an MOI of 0.1 then treated with 6-TG or 6-TG-Me. Supernatant was harvested at 24 hpi and stored at -80∞C until titering on BHK cells. (n=3 statistical significance was determined by paired ratio t test; *, p<0.05; **, p<0.01; ns, non-significant). **(C)** 293T cells were transfected with SARS-CoV-2 Spike or an empty vector control then treated with 6-TG, 6- TG-Me, or DMSO vehicle control. Lysates were harvested 24 h after transfection and probed by western blotting as indicated. **(D)** 293T cells were reversed transfected with siRNA and total RNA was harvested after 48 h. HPRT1mRNA abundance was measured by RT-qPCR (n=3, ±SEM statistical significance was determined by one way ANOVA; ***, p<0.001). **(E)** As in D but cells were infected with HCoV-OC43 at an MOI of 0.1 24 h after siRNA treatment and supernatant was harvested and titered as in **(B)** (n=4 ±SEM statistical significance was determined by paired ratio t test; **, p<0.01; ***, p<0.001; ns, non-significant).

### 6-Thioguanine inhibits Spike maturation by inhibiting an unidentified cellular GTPase

6- TG inhibits several different GTPases including Rac1, RhoA, and Cdc42 (37, 43, 44). This mode of action was previously shown to inhibit rotavirus replication at sub-micromolar concentrations (64). Here, we sought to determine if any of the known targets of 6-TG were responsible for defects in Spike maturation. We tested known inhibitors of Rac1 (Rac1 Inhibitor V), RhoA (Rhosin), and Cdc42 (CASIN) for cytotoxic effects in 293T cells after 24 h incubation using an alamarBlue assay to identify optimal sub-cytotoxic doses for further testing (**Figs. 6A, 6B)**. We also tested a GTPase agonist ML099 that can potently activate Rac1, RhoA, and Cdc42, among other GTPases (65). None of the three GTPase inhibitors had any effect on Spike processing or accumulation **(Fig. 6C)**. Because many additional GTPases have redox-sensitive cysteine residues that could interact with 6-TG, we reasoned that 6-TG could affect viral glycoproteins through an as-yet-unidentified GTPase. In support of this notion, we observed that the broadly acting small GTPase agonist ML099 maintained normal Spike processing in the presence of 6-TG **(Fig. 6D)**. Pre-treating the cells with ML099 before addition of 6-TG further enhanced this effect **(Fig. 6E)**. We attempted to rescue HCoV-OC43 titer in both HCT-8 and 293A infection models but instead found that ML099 was itself acting as an antiviral agent **(Fig. 6F)**. While this was worth testing, this result was unsurprising as viral infection likely requires specific or unperturbed GTPase function, including frequent cycling between GTP- and GDP-bound forms. Overall, these data suggest that 6-TG inhibits Spike maturation by inhibiting an unidentified cellular GTPase.

**Figure 6.**
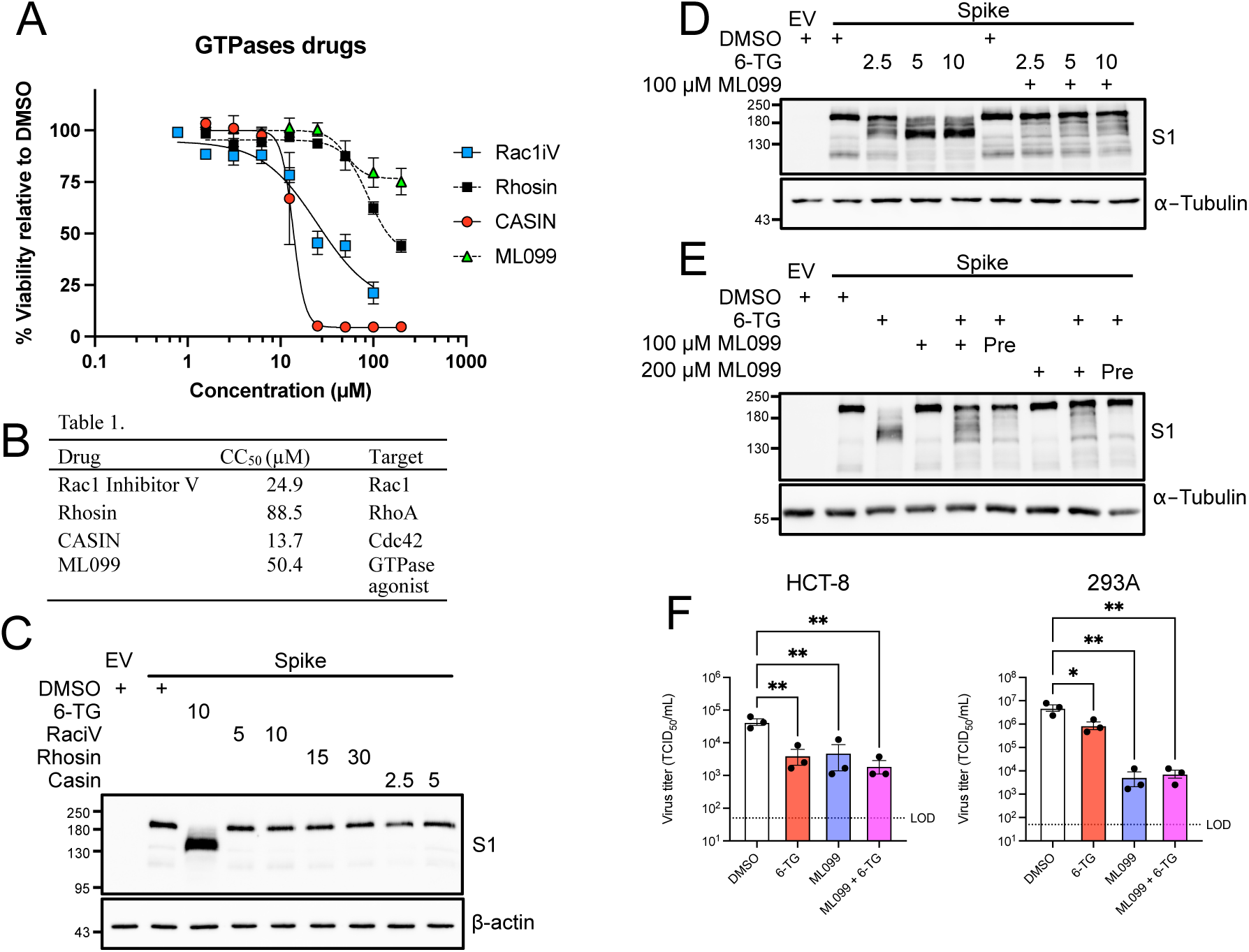
6-TG inhibits Spike maturation through GTPase inhibition. **(A)** alamarBlue cell viability assay of 293T cells treated Rac1 Inhibitor V (Rac1iV), Rhosin, CASIN, or ML099 (n=3±SEM). **(B)** Summary table of 50% Cytotoxic concentration (CC_50_) and inhibitor target of compounds tested in (A). **(C)** 293T cells were transfected with SARS-CoV-2 Spike or an empty vector control (EV) then treated with 6-TG, GTPase inhibitors, or DMSO vehicle control. Concentrations in legend are in µM. Lysates were harvested 24 h after transfection and were probed by western blotting as indicated. **(D)** 293T cells were transfected with plasmids encoding Spike or EV then co-treated with 6-TG, 100 µM ML099 GTPase agonist, or DMSO. Lysates were harvested 24 h after transfection and were probed by western blotting as indicated. Concentrations in legend are in µM. **(E)** as in (D) except some samples were pre-treated (Pre) for 4 h with ML099. **(F)** HCT-8 and 293A were infected with an MOI of ∼0.1 then treated with 10 µM 6-TG or DMSO vehicle control. After 24 h, the supernatants were harvested and stored at -80°C until titering on BHK-21 (n=3 ± SEM, statistical significance was determined by one-way ANOVA; *, p<0.05; **, p<0.01; ns, non-significant). LoD = Limit of Detection for virus titer.

## DISCUSSION

The reliance of viruses on the host cell biosynthetic and secretory apparatus, combined with the existence of varied and potent host antiviral mechanisms, makes host targeted antivirals (HTAs) useful additions to our armamentarium of direct-acting antivirals (DAAs). By targeting these host cell processes, HTAs may not be so easily subverted by the rapid viral evolution that frequently undermines DAAs. We recently reported that the thiopurines 6-TG and 6-TGo inhibit influenza A virus (IAV) by selectively disrupting the normal processing and accumulation of HA and NA glycoproteins (34). We hypothesized that other enveloped viruses that rapidly produce a burst of viral glycoproteins would also be susceptible to these thiopurines. Here, we report that the thioguanine 6-TG disrupts the normal processing and accumulation of CoV Spike glycoproteins and inhibits the release of infectious progeny. This drug-induced Spike processing defect disrupts the incorporation of SARS-CoV-2 Spike into VLPs or pseudotyped lentiviruses, which subsequently prevents infection of susceptible, ACE2-expressing cells. Thiopurines inhibit the small GTPase Rac1 by forming a covalent bond with a redox-sensitive cysteine, preventing further guanine nucleotide exchange. However, we show that selective chemical inhibitors of Rac1 or related small GTPases CDC42 and Rho had no effect on Spike processing and accumulation, whereas a broad GTPase agonist was able to counter the effects of 6-TG. This indicates that there are additional, unknown GTPase targets of 6-TG and that these are required for normal Spike processing and accumulation. Together, these findings suggest that functional small GTPases are essential for virus replication and that further development of GTPase inhibitors as host-directed antivirals is warranted.

We were intrigued by how 6-TG and 6-TGo strongly inhibited processing and accumulation of IAV and CoV glycoproteins. However, we also observed that these thiopurines had little effect on global host protein synthesis and secretion. Together, these observations suggest that viral glycoproteins may have properties that make them especially sensitive to these drugs. Certainly, viral glycoproteins must adhere to temporal or physical constraints that most host glycoproteins do not. All glycoproteins are synthesized, folded and post-translationally modified in the ER. However, viral glycoproteins can impose an acute burden on the ER when they are rapidly synthesized in large numbers in a relatively brief time frame. This burden may differ in severity depending on the extent of shutoff of host gene expression, as viruses with efficient host shutoff mechanisms would deplete host glycoproteins and facilitate viral utilization of the cellular biosynthetic and secretory apparatus. Viral envelope proteins are often more heavily glycosylated than their host counterparts, which can shield them from antibody recognition. Because N-glycosylation is a dynamic process involving glycan trimming and maturation, viral infection may impose specific burdens on these components of the ER quality control machinery. This may result in delayed trafficking of glycoproteins through the secretory pathway, leading to their degradation. Finally, it should be noted that viral glycoproteins are often assembled into large homo- and hetero-oligomeric complexes that must traffic to sites of assembly. Together, these aspects of viral glycoprotein biology place additional burdens on the ER and the associated machinery that governs glycoprotein folding, modification and trafficking.

Because cellular GTPases play important roles in viral glycoprotein biogenesis and trafficking, they may be good targets for host-directed antivirals. Viral assembly occurs at distinct sites like the plasma membrane (influenza A viruses) or the ERGIC (coronaviruses). Small molecules that disrupt efficient viral glycoprotein processing, oligomerization and trafficking could have the effect of ‘misplacing’ these important viral components and inhibiting essential steps in viral assembly and egress. We speculate that 6-TG and 6-TGo inhibit GTPases involved in key steps of viral glycoprotein synthesis, processing and trafficking, diverting them from their roles in viral assembly and consigning them to degradation via cellular catabolic processes. While these drugs are well-tolerated in humans, we believe that more can be gained by identifying new target GTPase(s) and further elucidating the antiviral mechanism of action to inform the development of novel chemical entities with superior properties.

## MATERIALS AND METHODS

### Cell lines

Human embryonic kidney 293T and 293A cells, human hepatoma Huh-7.5 (66) and African green monkey kidney Vero cells were all maintained in Dulbecco’s modified Eagle’s medium (DMEM; ThermoFisher, 11965118) supplemented with heat-inactivated 10% fetal bovine serum (FBS, ThermoFisher, A31607-01), 100 U/mL penicillin, 100 µg/mL streptomycin, and 2 mM L-glutamine (Pen/Strep/Gln; ThermoFisher, 15140122 and 25030081). HCT-8 cells were cultured as above with additional 1X MEM Non-Essential Amino Acids (NEAA, ThermoFisher) supplemented. Baby hamster kidney (BHK-21) were maintained as above yet supplemented with 5% FBS. African green monkey kidney Vero’76 and human lung adenocarcinoma Calu-3 cells were maintained in DMEM (Sigma-Aldrich, D5796; St. Louis, MO, USA) supplemented with 10% fetal bovine serum (ThermoFisher, 16000-044) and 1X Penicillin-Streptomycin (ThermoFisher, 15140148). hTert-immortalized human foreskin fibroblast (BJ) cells (a kind gift from William Hahn, Dana-Farber Cancer Institute, Boston, USA) cells were maintained in a 4:1 blend of knockout DMEM (ThermoFisher, 10829-018) and Medium 199 (ThermoFisher, 11150-059) with 15% FBS and Pen/Strep/Gln. All cells were maintained at 37°C in 5% CO_2_ atmosphere. To generate 293A-ACE2 cells, 293A cells were stably transduced with a lentivirus vector encoding ACE2 (pLJM1-ACE2-BSD) or an empty vector control (pLJM1-B*) (67), then selected and maintained in 10 µg/mL Blasticidin S HCl (ThermoFisher, A11139030).

### Chemical Inhibitors

6-thioguanine (6-TG, A4882), 2-amino-6-mercaptopurine riboside hydrate (6-TGo, 858412), 6-mercaptopurine (6-MP, 38171), brefeldin A (BFA; B7651), CASIN (SML1253), ML-099 (SML1258), Rac1 inhibitor V (5308200001), rhosin (555460), thapsigargin (Tg, cat#) and tunicamycin (Tm, cat#) were all purchased from Sigma-Aldrich. N9-methyl 6- thioguanine (6-TG-Me) was synthesized as described in Supporting Information. All drugs were solubilized in dimethyl sulfoxide (DMSO) and stored at -80°C. Stock concentrations were diluted to the indicated concentrations in cell culture media.

### Human coronavirus infections and quantitation

Stocks of severe acute respiratory syndrome coronavirus 2 (SARS-CoV-2) were propagated in Vero’76 cells (for strain SARS-CoV- 2/Canada/ON/VIDO-01/2020, used in Fig. 1A) or Vero E6 cells (SARS-CoV-2/SB3-TYAGNC, used in Figs. 3A, 3B). To generate virus stocks, cells were infected at a low MOI for 1 h in serum-free DMEM at 37°C following replacement of the inoculum with DMEM supplemented with 2% FBS and 1X Pen/Strep and continued incubation at 37°C. Once CPE reached 75% at 2-3 d post-infection, the viral supernatant was harvested, centrifuged at 5000 x *g* for 5 min, and then the cleared viral supernatant was aliquoted and stored at -80°C. Stocks were titered by plaque assay on Vero E6 cells (68) or TCID50 on Vero’76 cells with DMEM/2%FBS/Pen/Strep/Gln and calculated using the Spearman-Kärber method.

Stocks of human coronavirus OC43 (hCoV-OC43; ATCC, VR-1558) were propagated in Vero cells. Cells were infected at a MOI of 0.05 for 1 h at 33°C in serum-free DMEM. After 1 h, the infected cells were maintained in DMEM supplemented with 1% FBS and Pen/Strep/Gln for five days at 33°C. Upon harvest, the culture supernatant was centrifuged at 1000 x *g* for 5 min at 4°C, aliquoted, and stored at -80°C. Stocks of human coronavirus 229E (hCoV-229E; ATCC, VR- 740) were prepared similarly to hCoV-OC43 yet using Huh-7.5 cells and DMEM supplemented with 2.5% FBS and Pen/Strep/Gln. Viral titers were enumerated by median tissue culture infectious dose (TCID_50_) assays. Following the serial dilution of the samples, the appropriate cell line was infected for 1 h at 37°C prior to the replacement of the inoculum for the indicated overlay medium and incubated at 37°C: BHK-21 cells (hCoV-OC43, DMEM/1%FBS/Pen/Strep/Gln) or Huh-7.5 cells (hCoV-229E, DMEM/2.5%FBS/Pen/Strep/Gln). Viral titers were calculated using the Spearman-Kärber method.

To test the effect of the indicated compounds on hCoV replication, 293A, 293T, BJ, Calu- 3, HCT-8, or Huh-7.5 cells were infected with hCoV diluted in serum-free DMEM at the indicated MOI for 1 h. After 1 h, the inoculum was removed, replaced with DMEM/Pen/Strep/Gln containing 1% FBS and NEAA (HCT-8), 2% FBS (Calu-3), 2.5% FBS (Huh-7.5, 293A or 293T), or 5% FBS (BJ) supplemented with DMSO or drug at the indicated concentration, and maintained for an additional 23 h. Infected BJ cells were incubated at 33°C during virus replication; all other cells were incubated at 37°C. For infections of 293A, BJ, HCT-8, or Huh-7.5, the infected cells were scraped into the culture medium, subjected to three freeze/thaw cycles, then stored at -80°C. For SARS-CoV-2-infected Calu-3 cells or HCoV-OC43 infected 293T cells virus was collected from culture supernatants at 48 h post-infection and stored at -80°C. Titers of hCoV-OC43, hCoV- 229E, and SARS-CoV-2 were determined using TCID_50_ assays as indicated above.

### Cytotoxicity assays

Cells seeded into 96-well tissue culture plates were treated with drugs diluted at indicated concentrations and incubated with the cells for 20 h. At 20 h post-treatment, 10% alamarBlue Cell Viability Reagent (ThermoFisher Scientific, DAL1025) diluted in DMEM/Pen/Strep/Gln containing 1% FBS and NEAA (HCT-8), 2.5% FBS (Huh-7.5 or 293T), or 5% FBS (BJ) was added and further incubated for 4 h. Plates were read on FLUOstar Omega 96 well plate reader at an excitation of 544 nm, and an emission of 580-590 nm. Drug treatments were then normalized to DMSO. For viability experiments using Calu-3 cells, cells grown in 96-well plates were treated with drugs diluted at indicated concentrations and incubated with the cells for 48 h. At 48 h post-treatment, a one-fifth volume of CellTiter 96 AQueous One Solution Cell Proliferation Assay (G3580, Promega) was added to DMEM/Pen/Strep containing 2% FBS and incubated for 2 h. Plates were read at an absorbance of 490 nm with an xMark™ Microplate Absorbance Spectrophotometer (Bio-Rad). Values were imported into GraphPad PRISM (v.9) and CC_50_ values were calculated by fitting a non-linear curve to the data. Selectivity index (SI) was calculated by dividing the IC_50_ (calculated above) by the CC_50_.

### Real-time quantitative PCR

Total RNA from cells was extracted using the RNeasy Plus Mini Kit (QIAGEN, 74134) following the manufacturer’s protocol. Synthesis of cDNA was performed using the Maxima H Minus First Strand cDNA Synthesis Kit (ThermoFisher Scientific, K1652) using random primers. Quantitative PCRs were performed using 200 nM primers, 1:250 (final) diluted cDNA, and 1X GoTaq qPCR Master Mix (Promega, A6002). Amplifications were performed using a CFX Connect Real-Time PCR Detection System (Bio-Rad) and Bio-Rad CFX Manager 3.1 software. To measure supernatant virions, supernatant was mixed 1:1 with virus lysis buffer [20 mM Tris HCl pH7.4, 300 mM NaCl, 2.5% NP-40, and 40 U/mL RNAaseOUT (ThermoFisher)] for 5 min, then diluted 1:5 with nuclease-free water before 2 µL was used as template in a 10 µL GoTaq 1-Step RT-qPCR (Promega, A6020) reaction (69). Primer sequences are listed in Table 1.

**Table 1.**
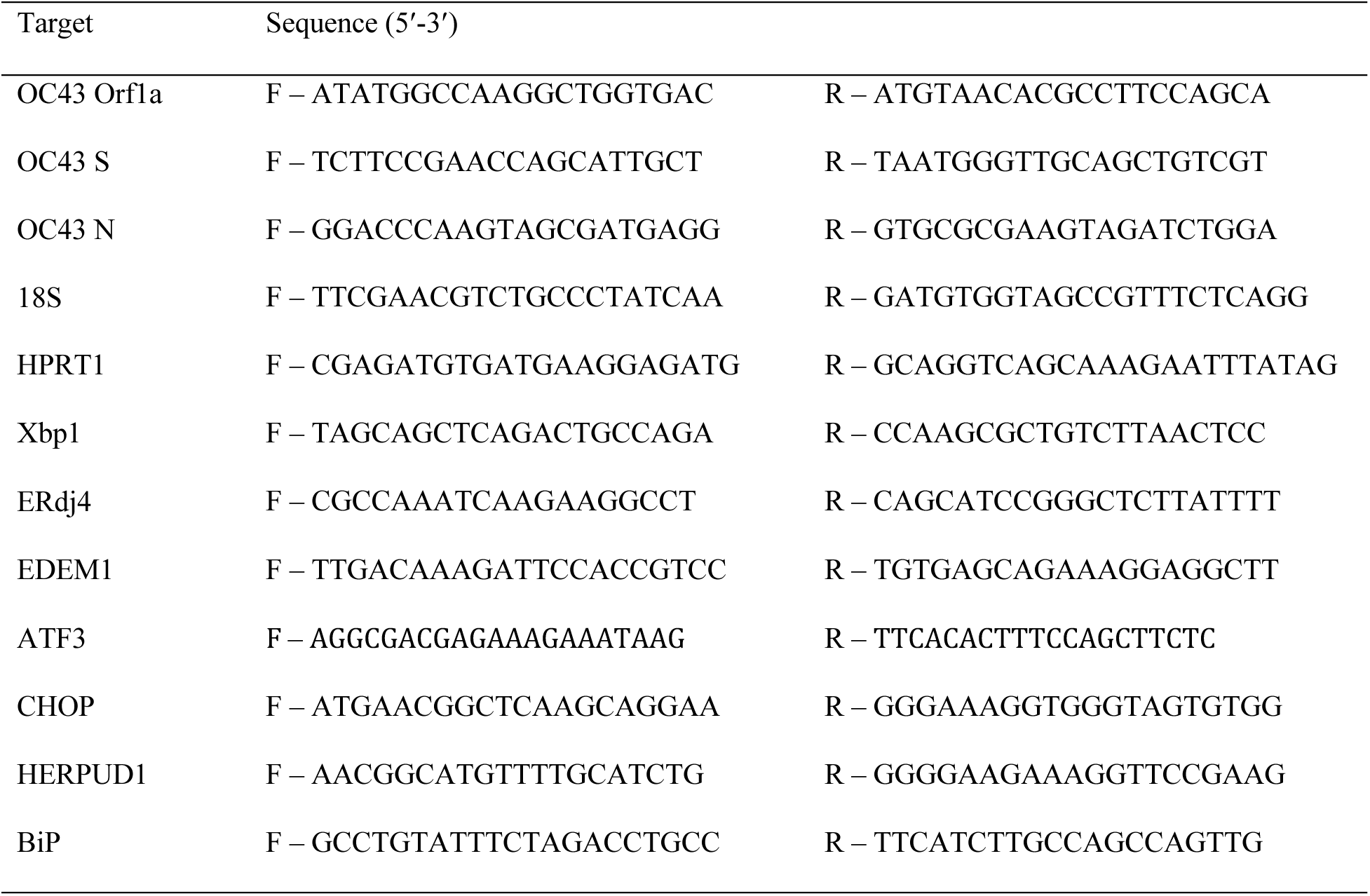
Primer sequences for RT-qPCR analysis.

### Immunofluorescence microscopy

293T were seeded on #1.5 coverslips (Zeiss, 411025-0003-000) coated with poly-L-lysine (Sigma, P2658). The following day the cells were infected with HCoV-OC43, to harvest the cells were washed once with 1x Dulbecco’s PBS (ThermoFisher) and fixed with 4% paraformaldehyde (Electron Microscopy Services, 15710) in PBS for 15 min at room temperature. Coverslips were blocked and permeablized in staining buffer [1% human serum (Sigma,4552) heat-inactivated at 56°C for 1h, 0.1% Triton X-100 PBS] for 1 h at room temperature, then stained with 1:500 dilution anti-OC43 N (MAB9012, Millipore Sigma) overnight at 4°C. The following day the coverslips were washed three times with PBS, then stained with goat anti-mouse Alexa 488 (ThermoFisher, R37120) in staining buffer for 1 h at room temperature in the dark. Coverslips were then counterstain for 5 min with Hoescht 33342 (ThermoFisher, 62249), washed three times with PBS, then mounted on coverglass using ProLong Gold anti-fade reagent (ThermoFisher, P36930). Z-stacks were imaged on a Zeiss LSM880 and processed into maximum intensity projections using Zen Black (Zeiss).

### Western Blotting

Cell monolayers were washed once with ice-cold PBS and lysed in 2x Laemmli buffer (4% [wt/vol] sodium dodecyl sulfate (SDS), 20% [vol/vol] glycerol, 120 mM Tris-HCl [pH 6.8]). DNA was sheared by repeated pipetting with a 21-gauge needle before adding 100 mM dithiothreitol (DTT) and boiling at 95^°^C for 5 min. Samples were stored at -20^°^C until analysis. Total protein concentration was determined by DC protein assay (Bio-Rad) and equal quantities were loaded in each SDS-PAGE gel. For ribopuromycylation assays, a modified version of the protocol by Schmidt, *et al.* (63) was used where the indicated samples were treated with 10 µg/ml puromycin (ThermoFisher, A1113803) for 10 min, followed by a PBS wash, and lysate preparation for immunoblotting as indicated above. Lysates from the ribopuromycylation assays were loaded into gels prepared using the TGX Stain-Free FastCast Acrylamide Kit (Bio-Rad, 1610183) (Fig. S1A). Lysates for quantitative Spike western blotting were loaded onto TGX Stain-Free 4-20% gradient gels (Bio-Rad, 4568094) (Fig. 3D). Peptide-*N*-Glycosidase F (PNGase F; NEB, P0704) treatment of lysates was performed as described in the manufacturer’s directions. All proteins after SDS-PAGE were transferred to polyvinylidene difluoride (PVDF) membranes (Bio-Rad) with the Trans-Blot Turbo transfer apparatus (Bio-Rad). Membranes were blocked with 5% bovine serum albumin in tris-buffered saline/0.1% [vol/vol] tween-20 (TBS-T) before probing overnight at 4^°^C with antibodies raised to the following targets: mouse anti-puromycin (Millipore-Sigma, MABE343), mouse anti-OC43 N (CoV antibody, OC43 strain, clone 541-8F, Millipore-Sigma, MAB9012), rabbit anti-OC43 S (Cusabio, CSB-PA336163EA-1HIY), rabbit anti-BiP (Cell signaling Technologies (CST), mouse anti-XBP1s (CST, #12782), mouse anti-CHOP (CST, #2895), rabbit anti-α-Tubulin (CST, #2125), rabbit anti-SARS-CoV-2 S1 RBD (Elabscience, E-AB-V1006), anti-SARS-CoV-2 E (abbexa, abx226552), anti-SARS-CoV-2 M (Novus Biologicals, NBP3-05698), rabbit anti-SARS-CoV-2 N (Novus Biologicals, NBP3-05730), mouse anti-FLAG (CST, #2368), mouse anti-HIV-p24 (Abcam, ab9071) anti-Strep tag (IBA, 2-1507-001) and anti- β-actin (CST, #4967). Membranes were washed with TBS-T and incubated with HRP-linked secondary antibodies prior to detection with Clarity ECL chemiluminescence reagent (Bio-Rad). All blots were imaged on a Bio-Rad ChemiDoc-Touch system. Molecular weights were determined using the Broad Range, Color Prestained Protein Standard (NEB, P7719) or Precision Plus Protein All Blue Prestained Protein Standards (BioRad, 1610373). Molecular weights in kDa are indicated on the left of blot images. For quantitation of western blots, the total antibody signal intensity was divided by the Stain-Free total protein signal. Images were analyzed using Image Lab 6.1 (Bio-Rad). Images were cropped and annotated using Affinity Designer (Serif).

### Plasmid generation and transfections

All plasmids were purified using the QIAprep Spin Miniprep or QIAfilter Plasmid Midi Kits (QIAGEN) and all restriction enzymes were purchased from New England Biolabs (NEB). The pcDNA3-SARS-CoV-2-Spike plasmid contains a codon-optimized ORF for Spike from GenBank NC_045512 that was synthesized by GenScript (a kind gift from David Kelvin) then cloned between the KpnI and BamHI sites of pcDNA3.1(+). A 19 residue C-terminal truncation of Spike (SΔ19) and Spike with a C-terminal FLAG tag was generated by PCR. A Spike mutant lacking the Furin cleavage site between S1 and S2 (P681S, R682G, R683G, R685del), and two-proline stabilizing mutants, K986P and V987P (S-2P) (70–72) was generated by PCR mutagenesis using KOD Xtreme Hot Start DNA Polymerase (Sigma, 71975-M). pcDNA3-Membrane (M) and pcDNA3-Envelope (E) were cloned from pLVX- EF1alpha-SARS-CoV-2-M-2xStrep-IRES-Puro and pLVX-EF1alpha-SARS-CoV-2-E-2xStrep- IRES-Puro (kind gifts from Nevan Krogan, available from Addgene #141386 and #141385) using PCR to remove the 2xStrep epitope tag. pCMV3-MERS-FLAG was purchased from SinoBiologicals (VG40069-CF). pLJM1-Luc2 used in generating Spike-pseudotyped lentiviruses was generated by cloning Luc2 from pGL4.26 (Promega) into pLJM1-B* (67). pLJM1-ACE2- BSD was cloned by from cDNA of generated from Caco-2 cells. Primer sequences are listed in Table 2. 293T cells were transfected using polyethylenimine (PEI; Linear, MW 25000 (Polysciences Inc., 23966) dissolved in water (pH 7.4). Both plasmids and PEI were diluted in OptiMEM (Gibco), then combined for 15 min before adding to cell monolayers in serum-free DMEM. PEI was used at 3:1 ratio to plasmid DNA. After 4 h, serum-free media was replaced with DMEM containing 10% FBS with indicated treatments unless otherwise indicated. For pre-treatments with ML099. the drug was applied at the time of transfection, then refreshed when the media was changed 4 h after transfection. Generally, cells were lysed for immunoblotting analysis at 24 h post-transfection (hpt). For flow-cytometry and immunofluorescence experiments, cells were instead transfected with FugeneHD (Promega, E2311) according to the manufacturer’s instructions.

**Table 2.**
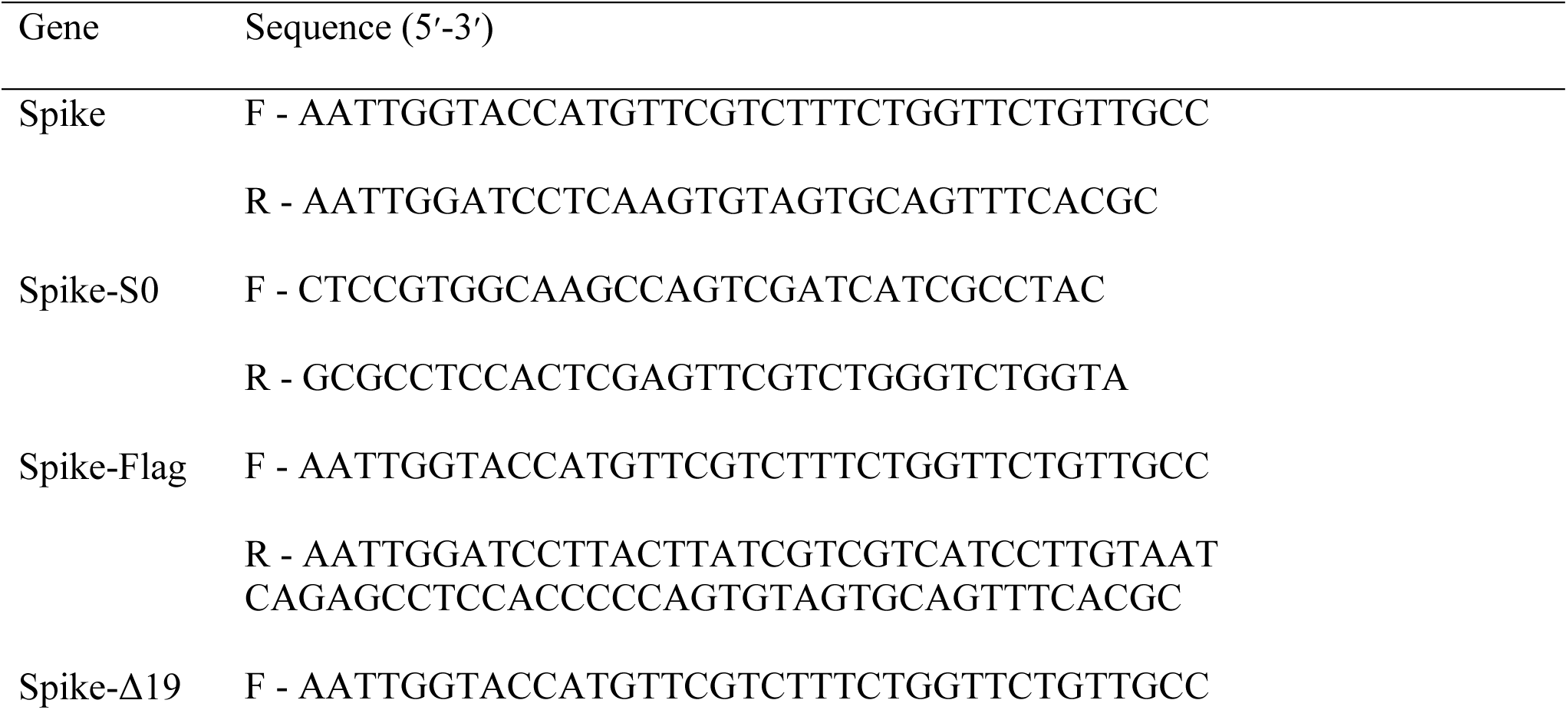
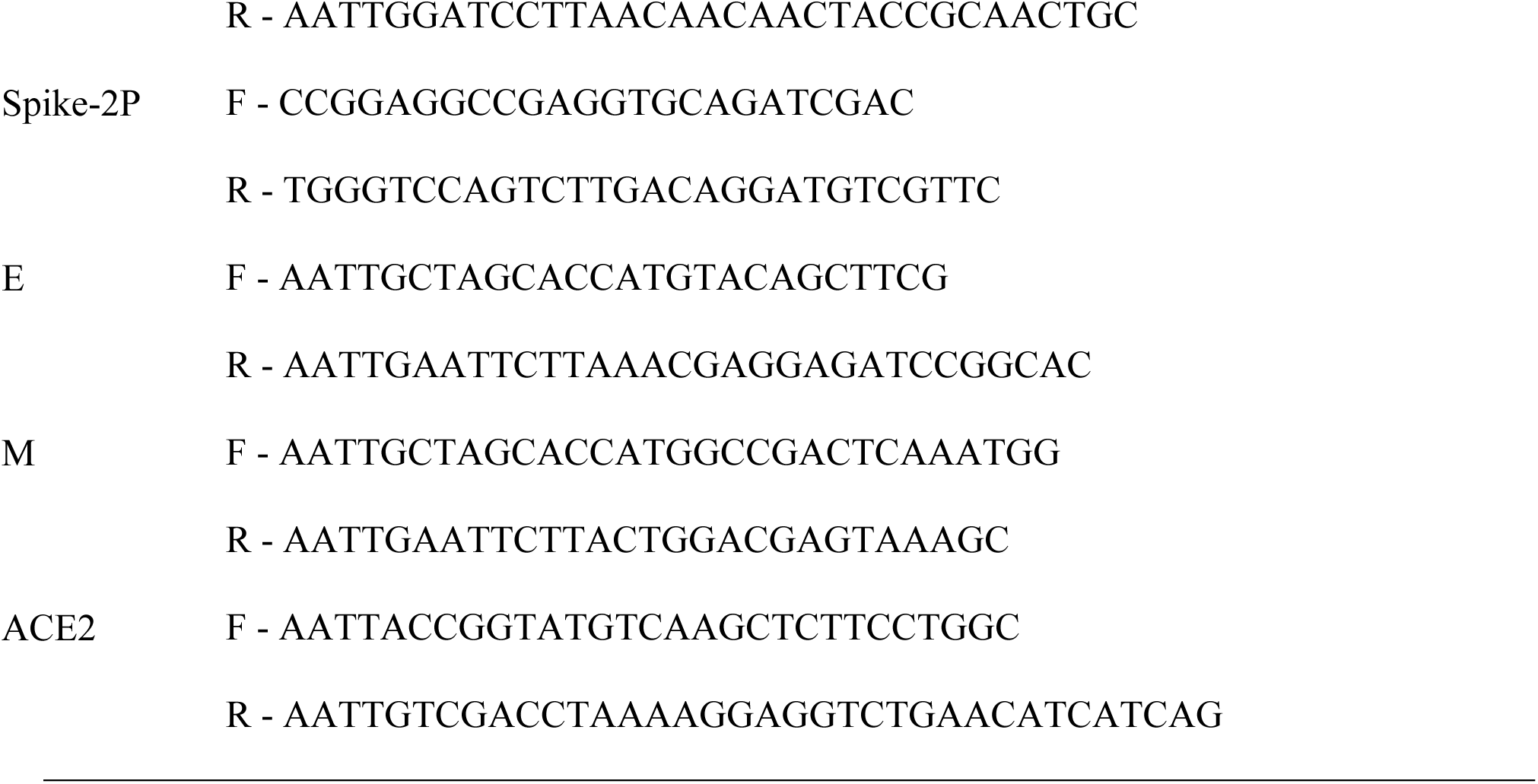
Primers used for cloning and PCR mutagenesis.

### siRNA knockdown experiments

All siRNAs were obtained from Invitrogen. 293T cells (1-1.5 x 10^5^ cells/mL) in 10% DMEM were reverse transfected with Lipofectamine RNAiMAX (Invitrogen, 13778075) and 10 nM of Silencer Select negative siRNA #2 (4390846), HPRT1 siRNA #1 (s6889), or HPRT1 siRNA #2 (s6888) diluted in OptiMEM (Gibco, 31985070). RNA was harvested from cells 48 hpt knockdown validation by RT-qPCR. For determining the effect of HPRT1 knockdown on viral titer after 6-TG treatment, the reverse transfected cells were infected at 24 hpt at an MOI of 0.1 then treated with DMSO or 10 µM 6-TG as described above. After a 24 h infection, the cell supernatant for titration, and the infected monoloayers were lysed with 2x Laemmli Buffer and processed for western blotting.

### Production of SARS-CoV-2 virus-like particles

To generate SARS-CoV-2 virus-like particles, 293T cells were seeded in a 10 cm dish and co-transfected with 2 µg of each pcDNA-Spike, pcDNA-M, pcDNA-E, and pLVX-EF1alpha-SARS-CoV-2-N-2xStrep-IRES-Puro (a kind gift from Neven Krogan, available from Addgene #141391) with PEI, then treated with 6-TG or DMSO as described above. After two days, supernatant was harvested and filtered through a 0.45 µm filter. For concentration and western blot analysis, filtered supernatant was underlaid with 20% sucrose PBS using a pipetting needle and centrifuged at 39 000 rpm for 2 h using an SW-41 rotor (Beckman Coulter). The supernatant and sucrose solution were carefully removed, and the pellet was resuspended in PBS, mixed in equal proportions with 2x Laemmli Buffer with 100 mM DTT and boiled at 95°C for 5 min before analysis by western blot.

### SARS-CoV-2 Spike-pseudotyped lentiviruses

To generate Spike-pseudotyped lentivirus particles, 293T cells were seeded in a 10 cm dish and co-transfected with 3 µg pLJM1-Luc2, 2 µg pSPAX2 (a kind gift from Didier Trono; Addgene #12260), and 2 µg of pcDNA3- SΔ19 with PEI, then treated with 6-TG or DMSO as described above. After two days, supernatant was harvested, filtered through a 0.45 µm filter, aliquoted and stored at -80°C until use. Genomes were quantified using a qPCR Lentivirus Titer Kit (Applied Biological Materials, LC900) as per the manufacturer’s instructions. Infectious pseudotyped particles were measured by infecting 293A cells stably expressing ACE2 or an EV control. Cell lysates were harvested 24 h after infection in Reporter Lysis Buffer (Promega, E397A), lysed by freeze thawing at -80°C, and luciferase activity was measured using Luciferase Assay reagent (Promega) on a FLUOstar Omega microplate reader (BMG Labtech). Lentiviruses that were concentrated and analyzed by western blot were prepared as described above for SARS-CoV-2 virus-like particles.

### Transmission electron microscopy (TEM)

Samples were prepared and visualized as described in (73). Briefly, HCoV-OC43 containing supernatant was concentrated by ultracentrifugation as described above, fixed with 2.5% glutaraldehyde (Electron Microscopy Sciences), then applied to Formvar/carbon coated grids and left to settle to 10 min. The grid was then rinsed with distilled water then stain with 2% w/v uranyl acetate for 30 s. Residual stain was wicked off with filter paper then left to air dry completely. Grids were imaged on a JEM 1230 TEM (JEOL) running at 80 kV using an OCRA-HR camera (Hamamatsu). Sample preparation, fixation, and TEM imaging were performed twice with representative images from one independent experiment shown.

### Quantitation of SARS-CoV-2 Spike surface expression

293T cells were co-transfected with pEGFP-N1 and plasmids expressing SARS-CoV-2 Spike (FL) or SARS-CoV-2 Spike-Δ19 for 24 h using FugeneHD (Promega). The cells were treated with DMSO or 10 µM 6-TG at 4 h post-transfection. At 24 h post-transfection, the cells were lifted with 10 mM EDTA PBS and pelleted at 300 x *g* for 5 min. Cells were washed once with wash buffer (2 mM EDTA, 2% FBS, PBS), then stained with 0.1 µg/sample Alexa Fluor 647-conjugated mouse-anti SARS-CoV-2 Spike S1 (R&D Systems, FAB105403R-100UG) at 4°C for 30 min in the dark. Cells were then washed prior to fixation with 2% paraformaldehyde (Electron Microscopy Sciences) in PBS at 4°C for 30 min in the dark, then washed and stored at 4°C in the dark until analysis using a FACSCelesta cell analyzer (BD Biosciences) configured with 405 nm, 488 nm, and 640 nm lasers using the same acquisition voltages for all replicates with at least 50000 single cells acquired per sample. The data were analyzed using FCS Express v. 6.06.0040 (De Novo Software) where transfected single cells were selected by EGFP fluorescence and analyzed for Spike surface expression via Alexa Fluor 647 fluorescence.

### *Gaussia* luciferase secretion assays

For short-term treatments, 293T were transfected with pCMV-*Gaussia* Luc Vector (ThermoFisher Scientific, 16147) as described above. After 18 h, medium was removed and replaced with fresh medium containing 6-TG, BFA, or DMSO. After 6 h medium was removed, debris was cleared by centrifuging samples at 5,000 *× g* for 5 min, then samples were stored at -80°C until measuring *Gaussia* luciferase activity using the *Gaussia* Luciferase Flash Assay Kit (ThermoFisher Scientific, 16158) as per the manufacturer’s directions on a FLUOstar Omega microplate reader (BMG Labtech). For long-term treatments, 293T cells were co-transfected with pCMV-*Gaussia* Luc and pLJM1-Luc2 and treated 6 h post transfection. After 24 h, supernatant harvested as described above. The cell monolayer was lysed in Reporter Lysis Buffer (Promega) as described above. To measure both Luciferase and *Gaussia* luciferase activity in samples, we used the Dual-Luciferase Reporter Assay System (Promega, E1910), which although designed for firefly and *Renilla* luciferase, can be readily adapted for firefly and *Gaussia* luciferase as both *Gaussia* and *Renilla* luciferase both use coelenterazine as a substrate.

### Statistical analysis

Statistical analysis and graphing were performed using GraphPad PRISM (v.8 or v.9). Statistically significant differences were determined using Dunnett’s multiple comparison test for one-way ANOVA and Šídák’s multiple comparison test for two-way ANOVA.

### Graphics

Molecular models were created with NCBI PubChem (http://pubchem.ncbi.nlm.nih.gov/) The model presented in Fig. 7 was adapted from “Coronavirus Replication Cycle”, by BioRender.com (2022). Retrieved from https://app.biorender.com/biorender-templates.

**Figure 7.**
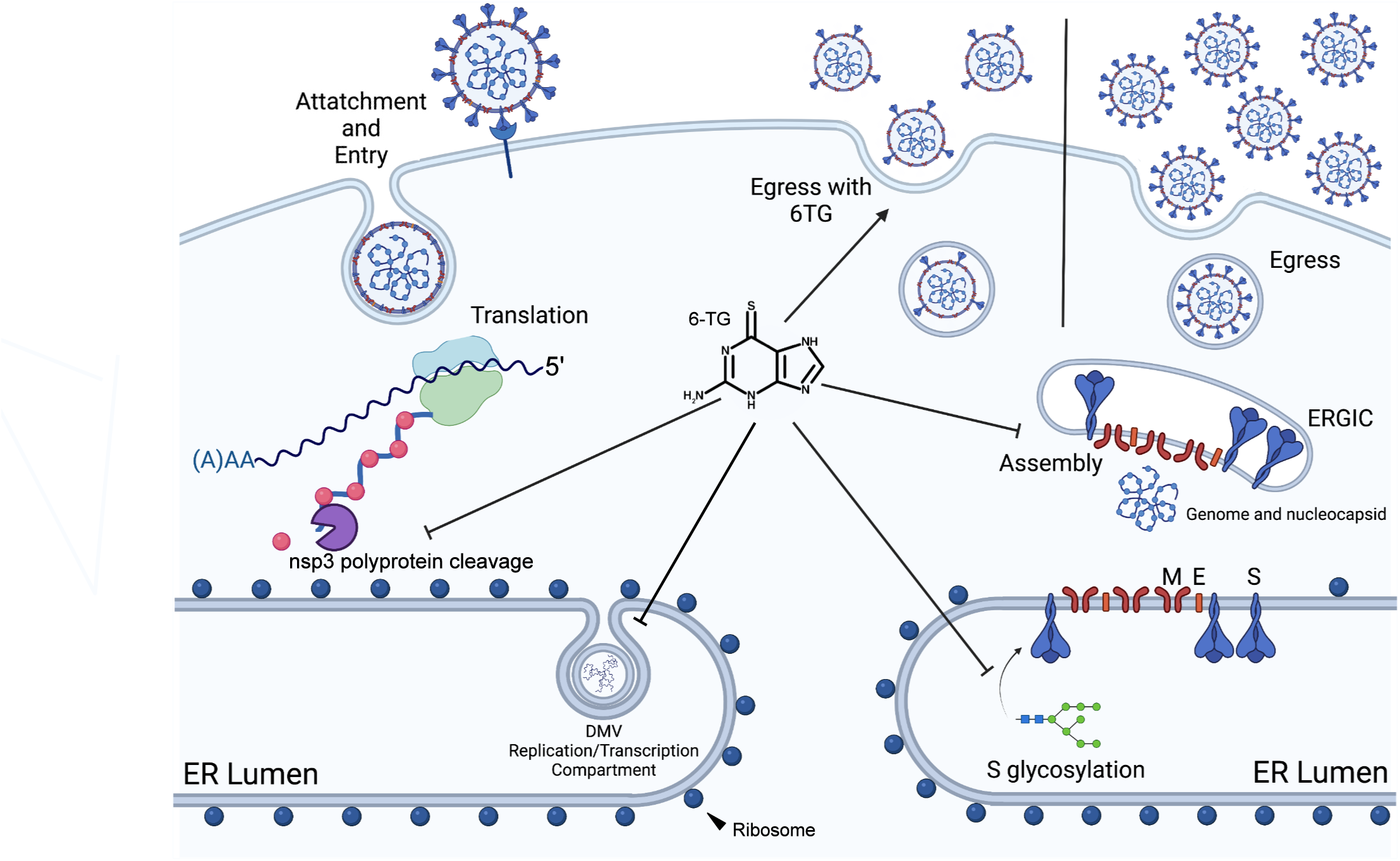
6-TG inhibits multiple essential steps in virus replication and assembly. 6-TG inhibits nsp3/papain-like protease (PLpro), delaying formation of replication compartments and reducing accumulation of viral RNA. 6-TG also inhibits normal glycosylation of Spike, which fails to be properly packaged in egressing virions. As a result, the total number of virions released from infected cells is reduced and those that are released have fewer Spikes on their surface.

## Supporting information

Supplemental Methods for chemical synthesis

## ACKNOWLEDGEMENTS

We are grateful for the support of the Dalhousie University Flow Cytometry Core Facility and the Cellular and Molecular Digital Imaging Core Facility. This work was supported by Canadian Institutes for Health Research (CIHR) Project Grant PJT-148727 (to C.M.), Natural Sciences and Engineering Research Council (NSERC) of Canada Discovery Grant RGPIN-2016-05083 (to S.L.B.), Coronavirus Variants Rapid Response Network (CoVaRR-Net) Grant 175622 (to J.A.C. and others), Research Nova Scotia Grant RNS-NHIG-2020-1383 and Lung Association of Nova Scotia Legacy Research Grant (to D.K.), and a Nova Scotia COVID-19 Health Research Coalition Grant to C.M. and E.S.P. SARS-CoV-2 research is supported in the laboratory of D.F. by the Canadian Institutes of Health Research (CIHR; OV5-170349, VRI-173022 and VS1- 175531). VIDO receives operational funding from the Government of Saskatchewan through Innovation Saskatchewan and the Ministry of Agriculture and from the Canada Foundation for Innovation through the Major Science Initiatives for its CL3 facility.

---

**Figure S1.**
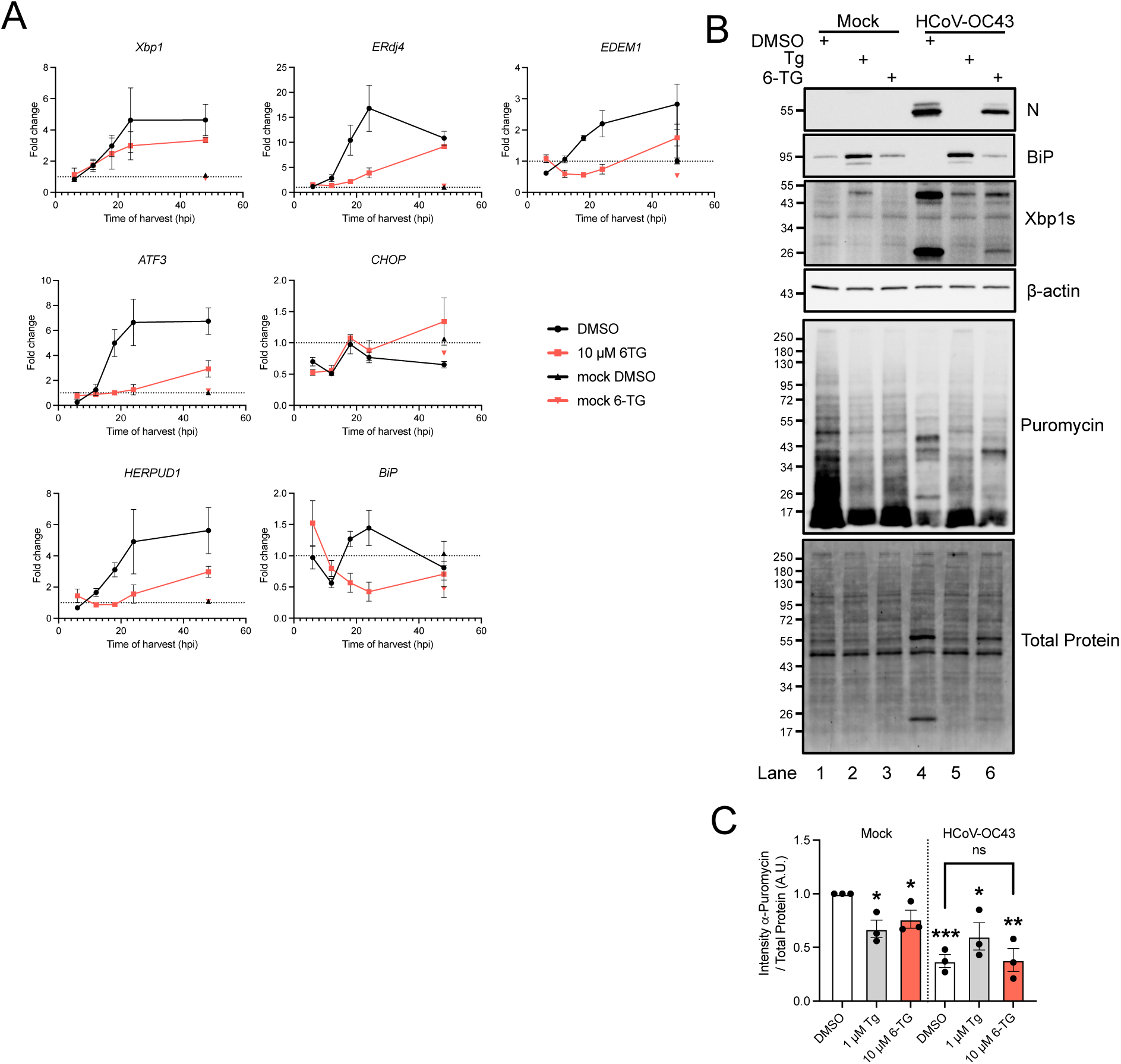
Activation of the unfolded protein response (UPR) during OC43 infection. **(A)** 293T cells were infected with HCoV-OC43 at an MOI of 0.1 then treated with 10 µM 6-TG or DMSO vehicle control. Total RNA from the cells were harvested at the times indicated and stored at -80∞C until RT-qPCR analysis of UPR target genes. **(B)** 293A cells were infected as in (A), then treated with 10 µg 6-TG, 1µg thapsigargin (Tg), or DMSO. Cells were treated with 10 µg/mL of puromycin for 10 min prior to harvest 24 h after infection. Lysates were probed by western blot as indicated. **(C)** Total puromycin intensity from (E) was quantified relative to total protein load and are graphed normalized to the mock-infected, DMSO-treated cells (n=3 ± SEM, statistical significance was determined by paired t-test compared to mock-infected, DMSO-treated cells; *, p<0.05; **, p<0.01; ***, p<0.001; ns, non-significant).

**Figure S2.**
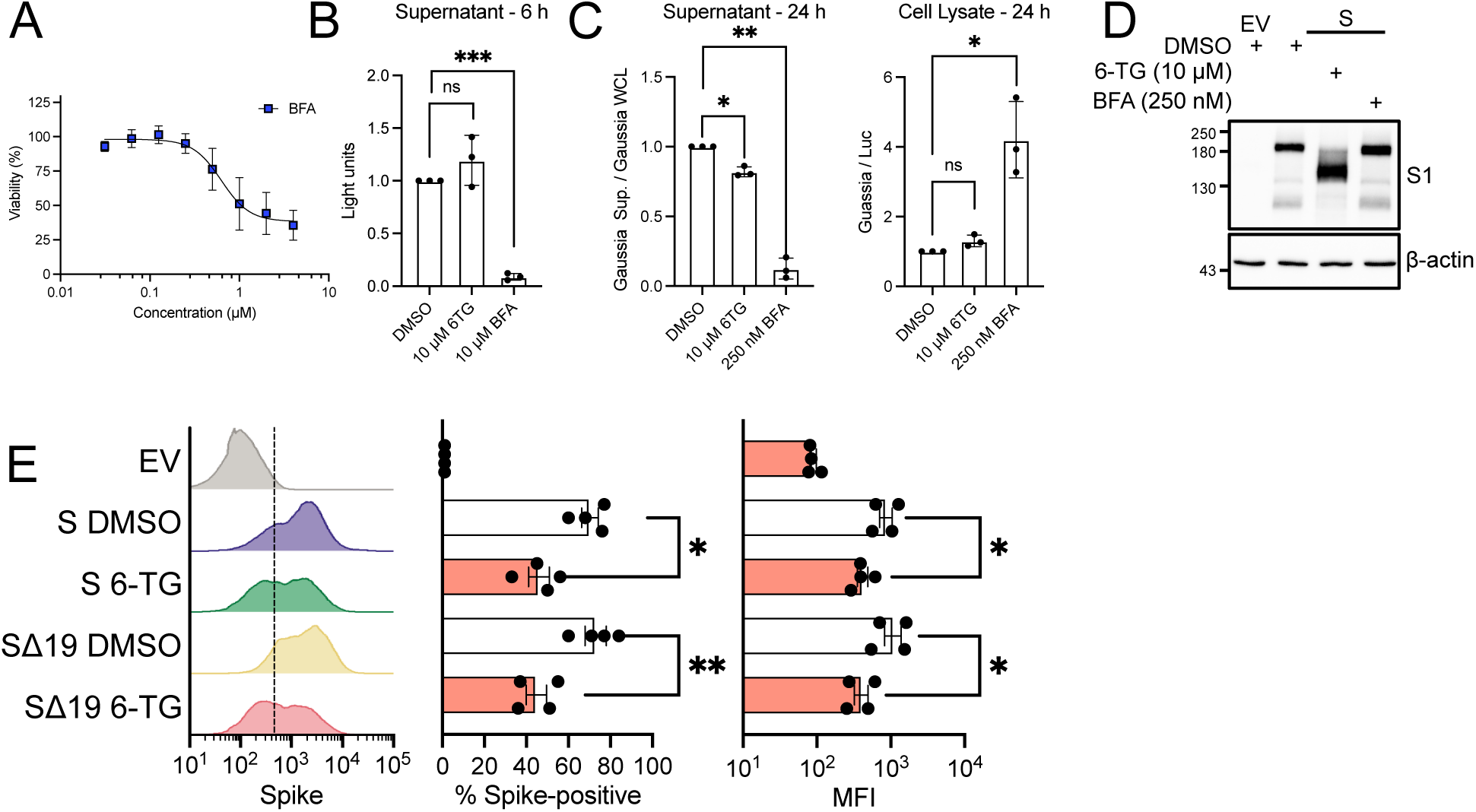
6-TG reduces surface expression of Spike but does not globally limit secretion. **(A)** alamarBlue cell viability assay of 293T cells treated with brefeldin A (BFA) (n=4±SEM). **(B)** 293T cells were transfected with *Gaussia* luciferase for 18 h then treated with 6-TG, BFA or DMSO vehicle control for 6 h. Supernatants were recovered from the cells and measured for luciferase activity. **(C)** 293T cells were transfected with both luciferase and *Gaussia* luciferase then treated with 6-TG, BFA, or DMSO. After 24 h, supernatants were removed and analyzed as in (B). Cell lysate was harvested in Reporter Lysis Buffer and stored at -80∞C until both luciferase and *Gaussia* luciferase activity was measured. (For B and C, n=3 ± SEM, statistical significance was determined by paired t-test compared to DMSO-treated cells; *, p<0.05; **, p<0.01; ***, p<0.001; ns, non-significant). **(D)** 293T cells were transfected with SARS-CoV-2 Spike or EV then treated with 6-TG, BFA, or DMSO. Lysates were harvested 24 h after transfection in 2x Laemmli buffer and were probed by western blotting as indicated. **(E)** 293T cells were co-transfected with EGFP and either Spike (S), Spike with 19 residue C-terminal truncation (SΔ19), or EV and then treated with 10 µM 6-TG or DMSO. After 24 h, cells were harvested, surface-stained for Spike then fixed prior to analysis by flow cytometry. EGFP+ cells were gated for analysis of number of Spike+ cell s and Median Fluorescent Intensity (MFI) (n=4±SEM statistical significance was determined by paired t-test between 6-TG and DMSO treated cells; *, p<0.05).

