## Supplemental Methods for chemical synthesis for "Thiopurines inhibit coronavirus Spike protein processing and incorporation into progeny virions"

**TABLE OF CONTENTS** **PAGE**


| | | |
|---|---|---|
| Materials and Methods | | S-3 |
| Figure S1: | $^{1}H$ NMR spectrum of 2-amino-9-methyl-3,9-dihydro-6H-purine-6-thione (**3**) | S-5 |
| Figure S2: | $^{13}C$ NMR spectrum of 2-amino-9-methyl-3,9-dihydro-6H-purine-6-thione (**3**) | S-6 |
| Figure S3: | ESI-HRMS spectrum of 2-amino-9-methyl-3,9-dihydro-6H-purine-6-thione (**3**) | S-7 |
| References | | S-8 |

### MATERIALS AND METHODS

**General.** 6-Chloropurine was purchased from Toronto Research Chemicals (Toronto, ON, Canada). Thiourea and all other chemicals were purchased from Sigma-Aldrich Canada Ltd. (Oakville, ON, Canada). All NMR spectra were obtained using a Bruker AVANCE 500 MHz spectrometer. Chemical shifts (*δ* in ppm) for proton ($^1$H) spectra are reported relative to the residual solvent signal for $CDCl_3$ (*δ* 7.26), DMSO-$d_6$ (*δ* 2.50), and HOD (*δ* 4.79).[1] Chemical shifts (*δ* in ppm) for carbon ($^{13}$C) spectra are reported relative to the residual solvent signal for $CDCl_3$ (*δ* 77.16) and DMSO-$d_6$ (*δ* 39.52).[1] Abbreviations in NMR spectra are: bm, broad multiplet; bs, broad singlet; bt, broad triplet; d, doublet; dd, doublet of doublets; m, multiplet; q, quartet; s, singlet; and t, triplet. High resolution (HR) electrospray ionization (ESI) mass spectra (MS) were collected using a Bruker microTOF Focus orthogonal ESI-TOF mass spectrometer instrument operating in either negative or positive ion mode. Melting points are uncorrected.

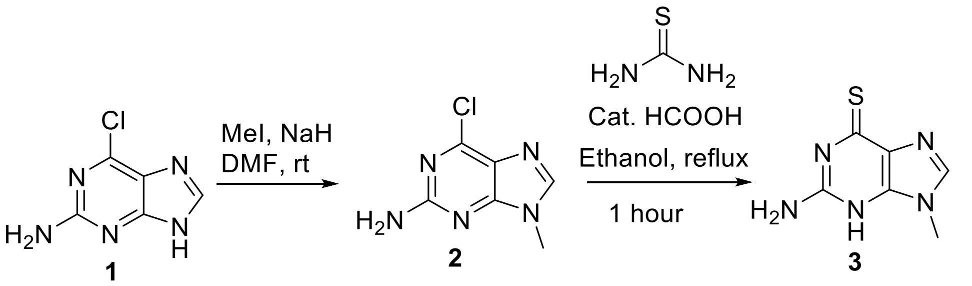


**6-Chloro-9-methyl-9H-purin-2-amine (2).** Following a procedure adapted from published protocols,[2,3] 6-chloropurine-2-amine (2.0 g, 11.8 mmol) was dissolved in of dry DMF (100 mL). NaH (0.56g, 60 wt.%, 14 mmol) was then added and the suspension was stirred for 15 min before addition of iodomethane (13 mmol). The reaction mixture was stirred at room temperature for 2 h, then the solvent was evaporated *in vacuo*. The residue was dissolved in ethyl acetate and the solution was subsequently washed with water, dried over anhydrous $MgSO_4$, and the solvent was evaporated *in vacuo*. The product was purified by silica gel chromatography (MeOH/$CHCl_3$; 1:99)

to afford 1.2 g of **2**. (56%). The $^{1}H$ and $^{13}C$ NMR spectra, as well as the mass spectrum, was in agreement with the literature.[2,3]

**2-amino-9-methyl-3,9-dihydro-6H-purine-6-thione (3).** Compound **2** (0.15 g, 0.82 mmol) was dissolved in ethanol (10 mL) followed by thiourea (0.25 gr, 3.27 mmol) and 2 drops of formic acid. The reaction mixture was the heated under reflux for 1 h. Upon cooling the reaction mixture to room temperature, a white precipitate formed, which was collected by filtration and washed twice with absolute ethanol (2 × 10 mL) to afford **3** as a white solid (120 mg, 81%); mp: >300 °C; $^{1}H$ NMR (500 MHz, DMSO) $\delta$ 12.46 (s, N$\underline{H}$, 1H), 8.56 (s, C$\underline{H}$, 1H), 7.26 (s, N$\underline{H}_2$, 2H), 3.63 (s, C$\underline{H}_3$, 3H); $^{13}C$ NMR (126 MHz, DMSO) $\delta$ 173.28, 153.99, 147.09, 140.82, 123.44, 30.17; HR-ESIMS: *m*/*z* calcd for $C_6H_7N_5Na_1S$ $[M+Na]^+$: 204.0314, found 204.0320.

**Figure S1.** [1]H NMR spectrum of 2-amino-9-methyl-3,9-dihydro-6H-purine-6-thione (**3**)

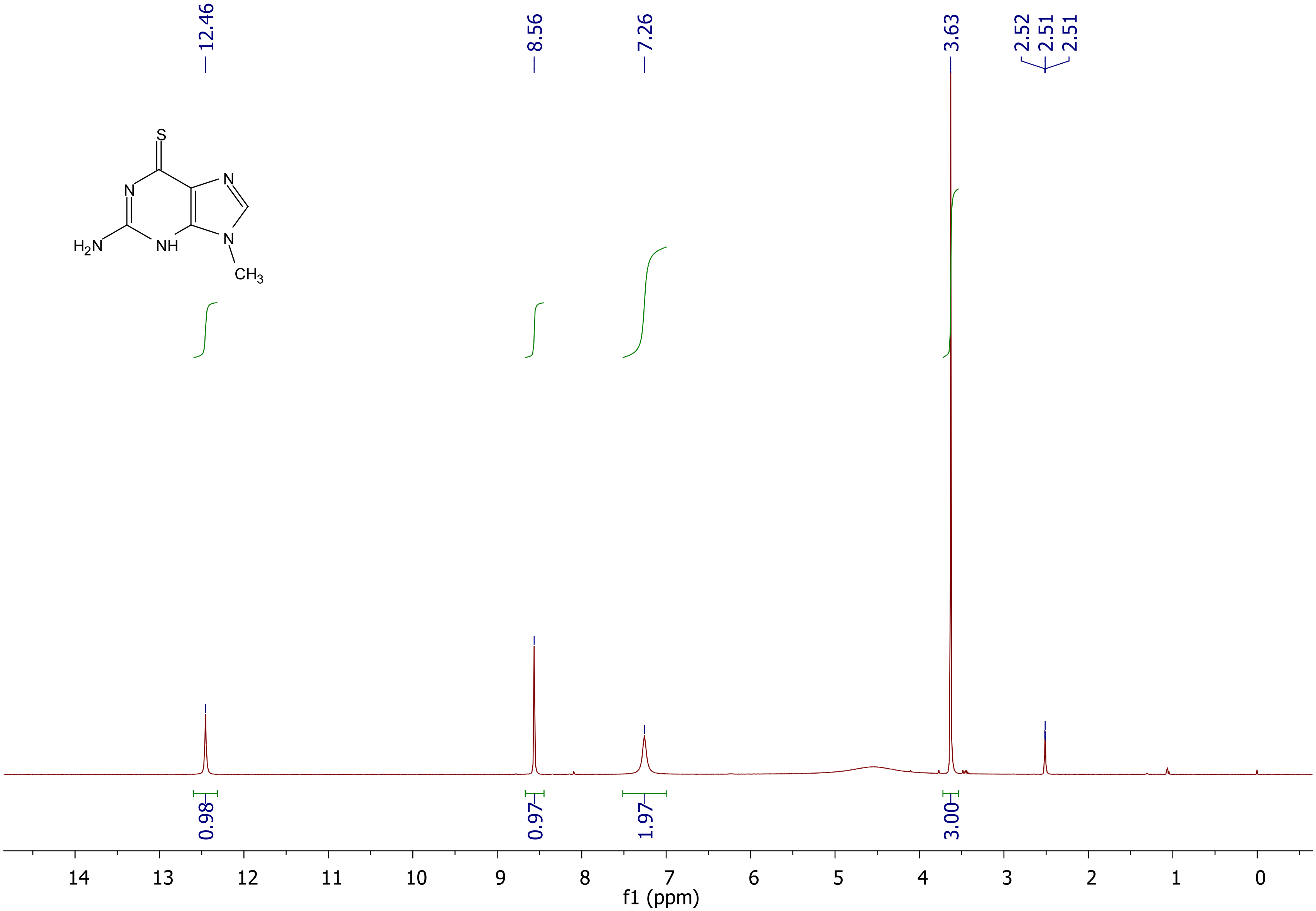

**Figure S2.** [13]C NMR spectrum of 2-amino-9-methyl-3,9-dihydro-6H-purine-6-thione (**3**)

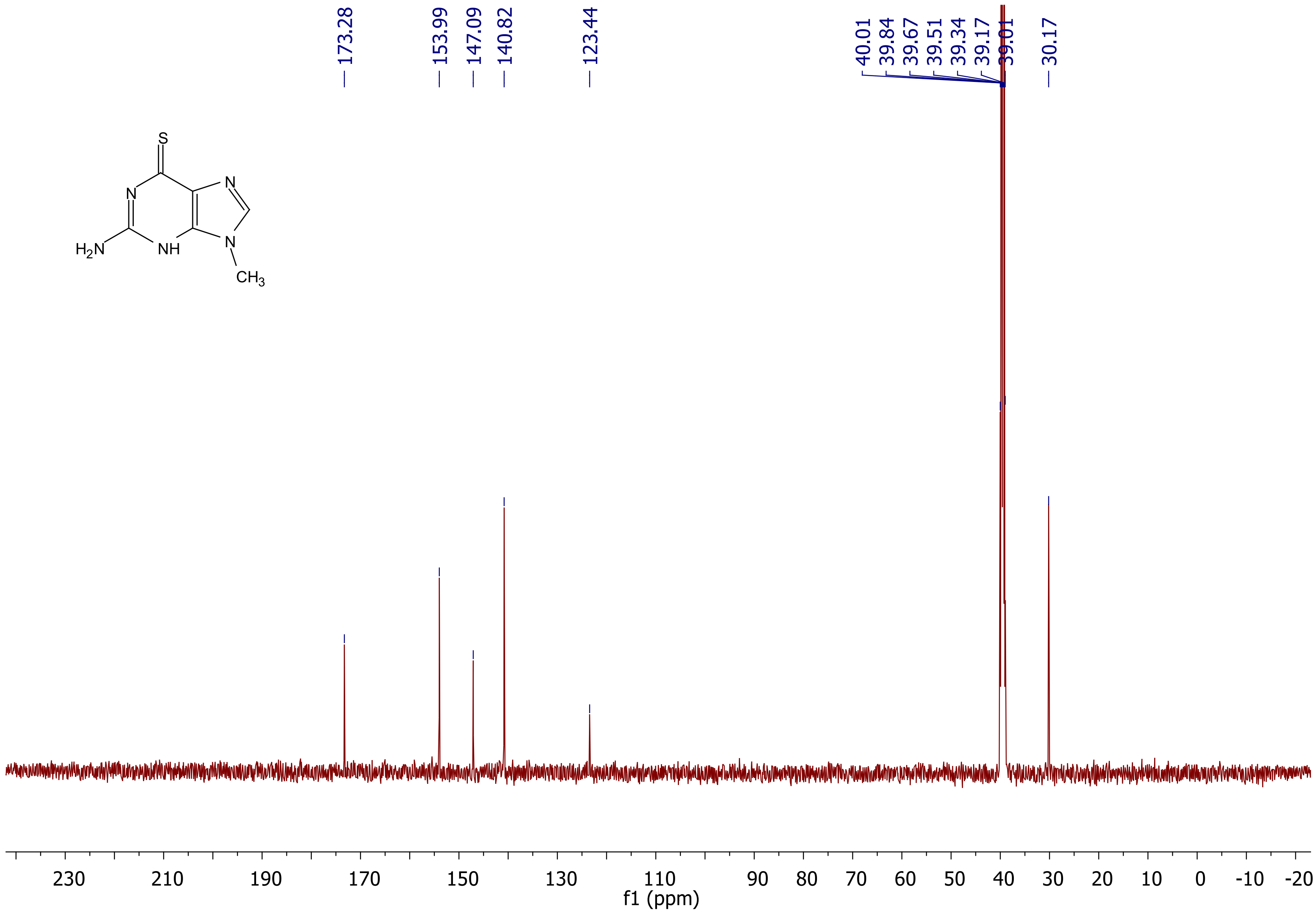

**Figure S3.** High resolution mass spectrum of 2-amino-9-methyl-3,9-dihydro-6H-purine-6-thione (**3**)

**Analysis Info**

| | | | | |
|---|---|---|---|---|
| Analysis Name | D:\Data\Xiao\Nov 15 2021000009.d | Acquisition Date | 11/15/2021 2:31:16 PM | |
| Method | Xiao all 1.m | Operator | x | |
| Sample Name | Methio-5B | Instrument | compact | 8255754.20059 |
| Comment | | | | |

**Acquisition Parameter**

| | | | | | |
|---|---|---|---|---|---|
| Source Type | ESI | Ion Polarity | Positive | Set Nebulizer | 1.0 Bar |
| Focus | Not active | Set Capillary | 3500 V | Set Dry Heater | 180 °C |
| Scan Begin | 50 m/z | Set End Plate Offset | -500 V | Set Dry Gas | 5.0 l/min |
| Scan End | 1500 m/z | Set Charging Voltage | 2000 V | Set Divert Valve | Source |
| | | Set Corona | 0 nA | Set APCI Heater | 0 °C |

| Meas. m/z | Ion Formula | m/z | err [ppm] |
|---|---|---|---|
| 204.031977 | C6H7N5NaS | 204.031437 | -2.6 |

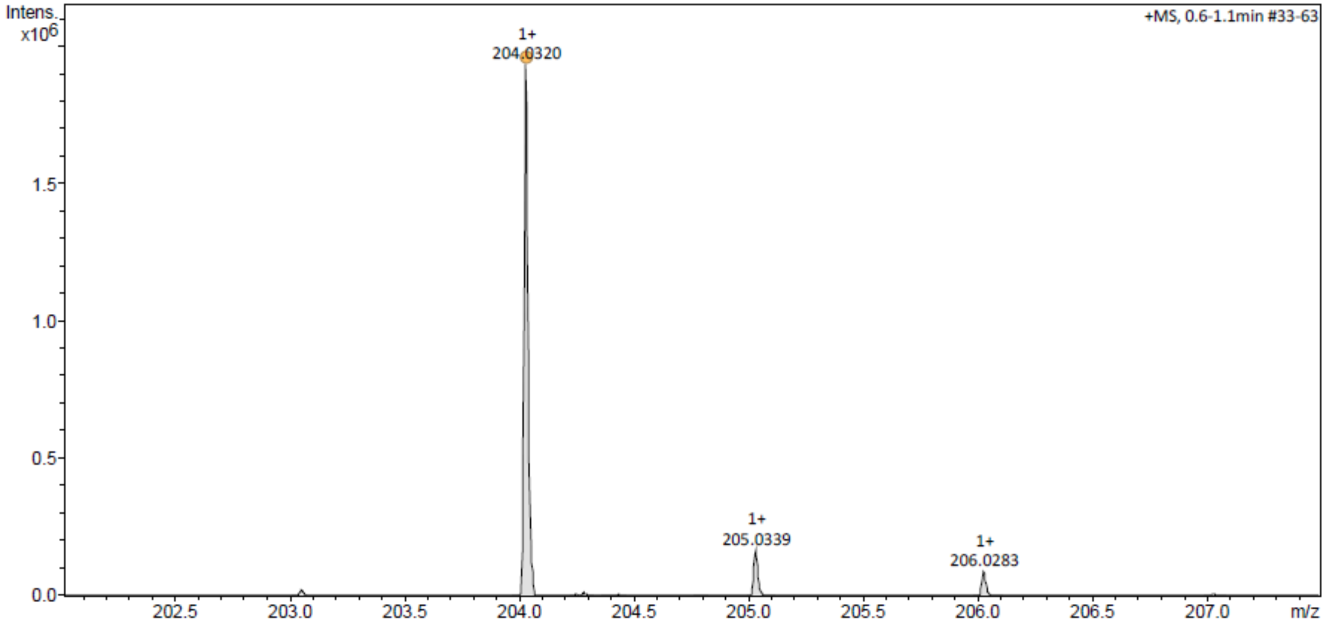


Nov 15 2021000009.d

Bruker Compass DataAnalysis 4.3 printed: 11/15/2021 2:36:23 PM by: x Page 1 of 1
